# Antigen presentation by discrete class I molecules on brain endothelium dynamically regulates T-cell mediated neuropathology in experimental cerebral malaria

**DOI:** 10.1101/2022.10.30.514412

**Authors:** CE Fain, J Zheng, F Jin, K Ayasoufi, Y Wu, MT Lilley, AR Dropik, DM Wolf, RC Rodriguez, A Aibaidula, ZP Tritz, SM Bouchal, LL Pewe, SL Urban, Y Chen, S Chang, MJ Hansen, JM Kachergus, J Shi, EA Thompson, JT Harty, IF Parney, J Sun, LJ Wu, AJ Johnson

**Author notes:** Correspondence to Aaron J. Johnson, PhD, Professor and Consultant, Departments of Immunology, Neurology, and Molecular Medicine, Guggenheim 4-11C, Mayo Clinic, 200 First St. SW, Rochester, MN 55905. **Author contributions:** CEF, JZ, FJ, KA, MTL, ARD, DMW, RCR, AA, and ZPT assisted with experiments. CEF, JZ, FJ, YW, SMB, YC, JMK, and JS analyzed data. CEF and AJJ designed experiments and interpreted results. CEF wrote the manuscript. All authors provided feedback for editing, read manuscript, and agree with data presented.

## Abstract

CD8 T cell engagement of brain vasculature is a putative mechanism of neuropathology in human cerebral malaria. To define contributions of brain endothelial cell MHC class I antigen-presentation to CD8 T cells in establishing this pathology, we developed novel H-2K^b^ LoxP and H-2D^b^ LoxP mice crossed with Cdh5-Cre mice to achieve targeted deletion of discrete class I molecules on brain endothelium. Using the *Plasmodium berghei* ANKA model of experimental cerebral malaria (ECM), we observe that H-2K^b^ and H-2D^b^ regulate distinct patterns of disease onset, CD8 T cell infiltration, targeted cell death, and regional blood-brain barrier (BBB) disruption. Strikingly, ablation of H-2K^b^ or H-2D^b^ from brain endothelial cells resulted in reduced CD8 T cell activation, attenuated T cell interaction with brain vasculature, lessened targeted cell death, preserved BBB integrity, and prevented ECM and the death of the animal. These data demonstrate that interactions of CD8 T cells with discrete MHC class I molecules on brain endothelium regulate development of ECM neuropathology. Therefore, targeting MHC class I interactions therapeutically may hold potential for treatment of cases of severe malaria.

## Introduction

*Plasmodium falciparum* has been classified as the deadliest parasite by the World Health Organization (WHO). This strain is also the most prevalent malaria parasite, accounting for majority of human infections^1^. Despite initiatives to limit mortality, it is estimated that greater than 600,000 people succumbed to malaria in 2020^2^. A severe complication *of Plasmodium falciparum* is severe encephalopathy, termed cerebral malaria (CM), which results in high mortality despite treatment^3, 4, 5, 6, 7^. CM accounts for approximately 80% of all malaria-related deaths, and patient admissions with CM are steadily increasing annually^2^. The majority of cerebral malaria patients are diagnosed by the presence of asexual parasite in a blood smear, deep coma, and absence of other infectious disease^3, 8^. Other hallmarks of disease can only be assessed post-mortem such as disruption of BBB tight junction proteins, vascular permeability, sequestration of parasitized red blood cell (prbc) and leukocytes, cerebral microhemorrhage and endothelial cell activation^9, 10^. MRI has been gaining traction as a diagnostic tool in patients, as vascular permeability can be assessed via T1 and T2 weighted MRI^10^. Recently, immunohistochemical studies of autopsied human brain specimens have revealed that CD8 T cells have been shown to be tightly associated with brain microvasculature and infiltrate brain parenchyma, implying an important role for this immune cell type in neuropathogenesis^11^.

*Plasmodium berghei ANKA* (PbA) infection is an established murine model that recapitulates human disease and CNS pathology, referred to as ECM^12^. As in the human disease, CD8 T cell responses are strongly implicated in the neuropathology. Importantly, CD8 T cells are required to have functioning cytotoxic capacity to develop the fatal BBB disruption associated with ECM^13, 14, 15, 16^. An IFN-y dependent upregulation of adhesion molecules (ICAM-1 and VCAM-1) and molecules associated with antigen (Ag) presentation (MHC Class I molecules D^b^ and K^b^ and MHC Class II) has been observed on brain and meningeal endothelial cells day 6 of PbA infection^17^. However, the contribution of specific cell types producing IFN-y in this pathology have not been interrogated^17^. Interest in these CD8 T cell-endothelial cell interactions, has prompted a concerted effort to define the requirement of Ag-specific CD8 T cells in this neuropathology. Due to the complex life-cycle of the parasite, defined by extensive antigenic variation throughout each stage, this has proven a difficult task. The engagement of CD8 T cells with class I molecules has been functionally assessed with bone marrow chimeras of MHC class I-deficient hosts reconstituted with class I-sufficient hematopoietic presentation^17^. However, it is known that CD8 T cells raised in these hosts have inherently impaired development and survival^17, 18^. While this important work narrows in on the significance of non-hematopoietic Ag-presentation in this pathology, the role of Ag-presentation specifically on brain vasculature has not yet been addressed. Furthermore, the contribution of discrete class I molecules in ECM pathology remain undefined.

In these studies, we sought to define the contribution of mouse brain microvascular endothelial cell (MBMEC) Ag-presentation by discrete class I molecules to the pathogenesis and lethality of ECM. We hypothesized that conditional ablation of Ag-presentation by H-2K^b^ and H-2D^b^ class I molecules on MBMECs would result in controlled modulation of activation and entry of parasite-specific CD8 T cells, attenuating pathology. To test this, we crossed endothelium-specific Cdh5-Cre mice with our novel H-2K^b^ LoxP and H-2D^b^ LoxP mice to create cKOs of discrete class I molecules. In these studies, we employed human and mouse transcriptomic data to evaluate gene expression in brains in response to plasmodium expression. We also employed *in vivo* 2-photon imaging to interrogate T cell interactions at the inflamed vasculature of the brain when individual MHC class I molecules are present or conditionally deleted on endothelium during PbA infection. Infiltration, activation, and localization of CD8 T cells and their potential targets were assessed via spectral flow cytometry in these novel transgenic mice. We demonstrate that interactions with specific class I molecules determined CD8 T cell activation, infiltration of the brain and localization and onset of cell death and neuropathology. However, knockout of H-2K^b^ and H-2D^b^ in endothelial cells prevented CD8 T cell interactions at brain vasculature, CD8 T cell activation, BBB disruption, cell death and neuropathology. These mice did not develop ECM, resulting in almost complete survival. Together these data demonstrate that CD8 T cells engage H-2K^b^ or H-2D^b^ class I molecules on MBMECs in order to coordinate activation, entry, and induction of neuropathology. These studies indicate that modulation of brain microvascular endothelial cell class I restricted Ag-presentation may hold therapeutic benefit in cases of cerebral malaria.

## Results

### Mouse and human brain microvascular endothelial cells increase antigen presentation molecules capable of eliciting T cell responses after *Plasmodium* infection

C57BL/6 mice were intraperitoneally (I.P.) injected with PbA prbcs, resulting in peripheral infection and manifestation of ECM on days 6-8 (Fig. 1a). Brains were harvested and RNA was extracted from whole brains of infected C57BL/6 mice and uninfected controls (Fig. 1a). Nanostring analysis was performed on bulk RNA for analysis of differential gene expression between uninfected and infected mice (Fig. 1b). Upregulated genes were associated with major immunological pathways, including Ag-presentation, T cell responses, cytotoxicity, cell activation and apoptosis. Downregulated genes were associated with markers of homeostasis or components of blood-brain barrier (Fig. 1b). Gene expression data were analyzed using Ingenuity Pathway Analysis (IPA). Causal network analysis identified MHC Class I family as the most significant master regulator in ECM pathology (Supplementary Table 1). MHC Class I family activation led to downstream activation of the majority of upregulated genes of interest identified in Fig. 1b (Fig. 1c). Canonical pathway analysis identified the most differentially enriched signaling pathways during ECM, confirming Ag-presentation as a major pathway of interest (Fig. 1d). MBMECs have been implicated to be involved in Ag-presentation during ECM (Fig. 1e). To interrogate their role, we employed enzymatic digestion and high-dimensional spectral flow cytometry to measure expression of Ag-presentation-associated molecules. MHC Class I median fluorescence intensity is increased on MBMECs during ECM and the percent of expression on MBMECs is significantly higher during ECM (Fig. 1f). MHC Class II is a well-known marker of antigen presenting cells (APCs). MHC Class II expression on MBMECs greatly increased from minimal to high during ECM, implicating their role in Ag-presentation (Fig. 1g). We next interrogated gene expression signatures of human brain microvascular endothelial cells (HBMECs) from human donor after exposure to *Plasmodium falciparum*. To this end, we utilized human microarray dataset from Tripathi et al. (accession GSE9861)^19^ in NCBI GEO database. In brief, isolated HBMECs were co-cultured with *Plasmodium falciparum* prbcs or uninfected red blood cells in culture and then analyzed (Fig. 1h). We first assessed the distribution of expression values for each human sample to ensure suitability for comparison (Supplementary Fig. 1a). After human sample validation we evaluated MHC Class I and MHC Class II expression levels. MHC Class I expression was significantly increased in HBMECs exposed to *Plasmodium falciparum* infected red blood cells (Fig. 1i). Human samples which were exposed to infection were subdivided into two groups 1) high adhesion molecule expression and 2) low adhesion molecule expression^19^ (Supplementary Fig. 1b). MHC Class II expression was shown to be significantly upregulated only in the high adhesion molecule samples compared to controls (Fig. 1j). Using GEO2R analysis platform, MHC Class I was identified as one of the most significantly differentially expressed genes and was in the top 5 genes with significant upregulation on HBMECs after exposure to *Plasmodium* infection (Supplementary Table 2). When comparing mouse and human genes involved with Ag-presentation we saw that gene activation and inhibition in these pathways were highly analogous (Supplementary Fig. 2). We also saw comparable upregulation of homologous genes of interest downstream of MHC Class I in human HBMECs, consistent with our observations in the mouse model (Fig. 1k). When performing gene set enrichment analysis (GSEA) on HBMEC transcriptomics data we identified the molecular signature for T cell activation involved in immune response to be highly enriched in HBMECs exposed to infection. The enrichment of this gene set indicates that HBMECs become an APC during *Plasmodium* infection, capable of eliciting this downstream T cell response (Fig. 1l). Together, these results suggest that in both mouse and human, brain endothelial cells are performing Ag-presentation to activate downstream T cell effector activities following *Plasmodium* infection.

**Fig 1.**
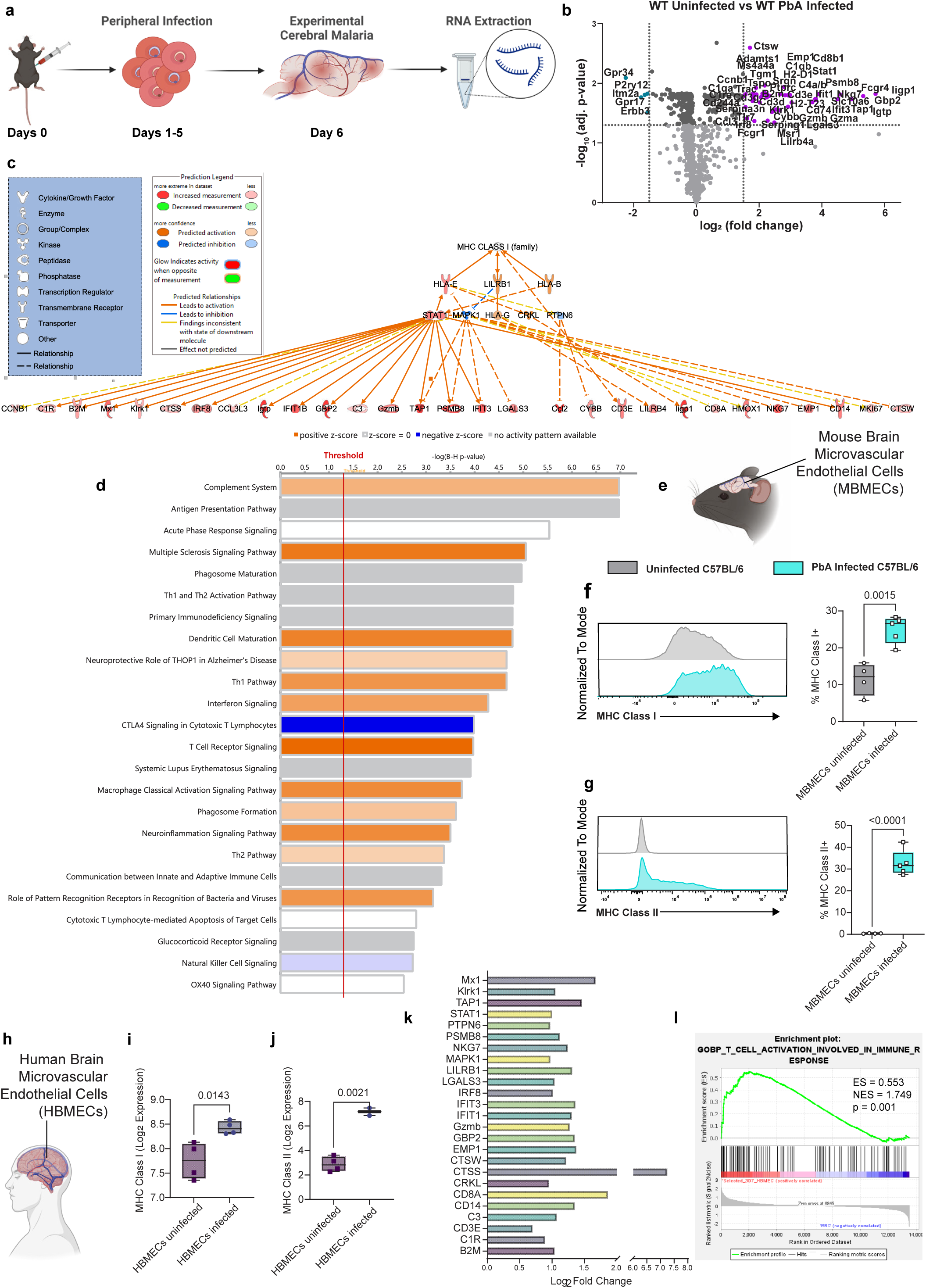
Mouse and human brain microvascular endothelial cells become antigen presenting cells during cerebral malaria. a) Flowchart showing steps of peripheral malaria infection, development of experimental cerebral malaria and RNA extraction for Nanostring analysis. b) Volcano plot of differentially expressed genes in whole brains of uninfected C57BL/6 mice versus PbA infected mice on day 6 of infection. Significantly upregulated genes are indicated on upper right (purple dots). Significantly downregulated genes are indicated on upper left (turquoise dots). Horizontal dotted line indicates cutoff of adjusted p-values (-Log10 of adjusted p-value 1.3 corresponding to a threshold of p-value of 0.05). Vertical dotted line indicates cutoff of log2 Fold change (-1.5 to 1.5). c) Ingenuity Pathway Analysis (IPA) was performed, and Causal network analysis identified MHC Class I family as the most significant network master regulator in experimental cerebral malaria pathology which led to the differential expression of the genes of interest. Depicted in c) a schematic of the homologous master regulator pathway and corresponding downstream genes. Coloring of the molecules corresponds to calculated z-scores (algorithm predicting activation or inhibition). Red indicates increased gene activation, orange indicates predicted activation of pathways, green indicates decreased gene activation, blue indicates predicted inhibition of pathways, white indicates a z-score of zero with equal evidence for activation and inhibition, grey indicates that z-score could not be calculated. d) Canonical pathway analysis identified significantly differential signaling pathways. P-value of overlap was adjusted using the Benjamini-Hochberg correction method. Bars are arranged in order of significance score with threshold (red line) set at p-value 0.05 (corresponding to a negative log value of 1.3). Coloring of bars corresponds to calculated z-scores (algorithm predicting activation or inhibition). Orange bars indicate activated pathways, blue indicates inhibited pathways, white indicates a z-score of zero with equal evidence for activation and inhibition, grey indicates that z-score could not be calculated. e) Schematic depicting C57BL/6 and naming convention of mouse brain microvascular endothelial cells (MBMECs). High-dimensional flow cytometry was performed on mouse brains on day 6 of PbA infection and uninfected age and sex matched controls. f) Representative histogram of MHC Class I expression on MBMECs and quantified frequency of MHC Class I high endothelial cells. g) Representative histogram of MHC Class II expression on MBMECs and quantified frequency. h) Schematic depicting human subjects and naming convention of human brain microvascular endothelial cells (HBMECs). i-j) Box plots depicting Log_2_ expression data from human endothelial cells in published microarray (GEO Accession: GSE9861). i) MHC Class I Log_2_ expression in control HBMECs vs. *Plasmodium* exposed HBMECs. j) MHC Class I Log_2_ expression in control HBMECs vs. *Plasmodium* exposed HBMECs selected as highly activated (with high expression of adhesion molecules). k) Log_2_ Fold change of genes of interest which lie downstream of the MHC class I signaling pathway in HBMECs during *Plasmodium* falciparum infection. GSEA was performed on HBMEC transcriptomic data. l) Enrichment plot showing enrichment of molecular signature for T cell activation involved in immune response pathway. The y-axis represents enrichment score (ES) and on the x-axis are genes (vertical black lines) represented in gene sets. The green line connects points of ES and genes. ES is the maximum deviation from zero as calculated for each gene going down the ranked list and represents the degree of over-representation of a gene set at the top or the bottom of the ranked gene list. The colored band at the bottom represents the degree of correlation of genes with the phenotype (red for positive and blue for negative correlation). Significance threshold set at nominal p-value < 0.05. Gene sets for T cell activation involved in immune response were significant only in HBMECs during infection. Data were analyzed with unpaired t test (f, g, i, and j). Box plots present the mean ± SD. All p-values are shown or ns for non-significant p-value *(p > 0.05)*.

### Endothelial cell specific K^b^ conditional knockout mice develop normal peripheral *Plasmodium* infection and immune responses

We crossed (class I deficient background, K^b^−/−, D^b^−/−) K^b^ LoxP mice with (class I deficient background, K^b^−/−, D^b^−/−) Cdh5-Cre mice (Fig. 2a). The resulting animals are defined as K^b^ Cre- littermate (with ubiquitous K^b^ expression), and Cdh5-Cre K^b^ cKO (with endothelial-specific ablation of K^b^), respectively. At baseline, Cdh5-Cre K^b^ cKO mice had significantly reduced protein levels of K^b^ on brain endothelium, as well as reduced percentage of brain endothelial cells that express K^b^ (Fig. 2b-d). T cell development was intact in Cdh5-Cre K^b^ cKO and K^b^ Cre- littermates with no changes in thymic T cell subset frequencies, mTEC and cTEC frequencies, or thymic weights between groups (Fig. 2e-f and Supplementary Fig. 3a). Furthermore, no sex specific differences pertaining to T cell development, overt changes in CD8 T cell repertoire, or change in splenic weights were observed (Supplementary Fig. 3b-d). To determine the effect of K^b^ class I molecule ablation on MBMECs during ECM, adult Cdh5-Cre K^b^ cKO (∼3-month-old) mice and K^b^ Cre- littermates were I.P. infected with 17#x00D7;10^6^ PbA prbcs and tissues were harvested at approximately 6-8 days post infection (dpi) (Fig. 2g). During infection K^b^ remained significantly reduced on brain endothelium (Fig. 2h-j). In the spleen, we observed no changes to granulocyte or APC subsets during infection in Kb Cre- littermates or Cdh5-Cre K^b^ cKO mice (Fig. 2k-l). Importantly, there were no differences in the ability to establish blood stage infection in these mice (Fig. 2m). Populations of immune cells were assessed in peripheral tissues at baseline and during infection and showed no changes in CD45+ immune cell numbers between Cre- littermates and Cdh5-Cre K^b^ cKO spleen, peripheral blood or inguinal LN (Supplementary Fig. 3e). In these peripheral tissues CD4 and CD8 T cell numbers and activation state remained unchanged between groups in all peripheral organs, with the exception of a modest reduction in numbers of activated CD8s at baseline in inguinal LNs in the Cdh5-Cre K^b^ cKO mice (Supplementary Fig. 3g). However, during PbA infection, there were no differences in CD8 activation in inguinal LNs (Fig. 2o). These results demonstrate that our Cdh5-Cre K^b^ cKO mouse model has efficient deletion of K^b^ on vasculature, while maintaining a fully functional peripheral immune response to *Plasmodium* infection. This allows us to proceed with interrogation of potentially brain-specific immunomodulatory functions of K^b^ Ag-presentation by brain endothelial cells in ECM.

**Fig 2.**
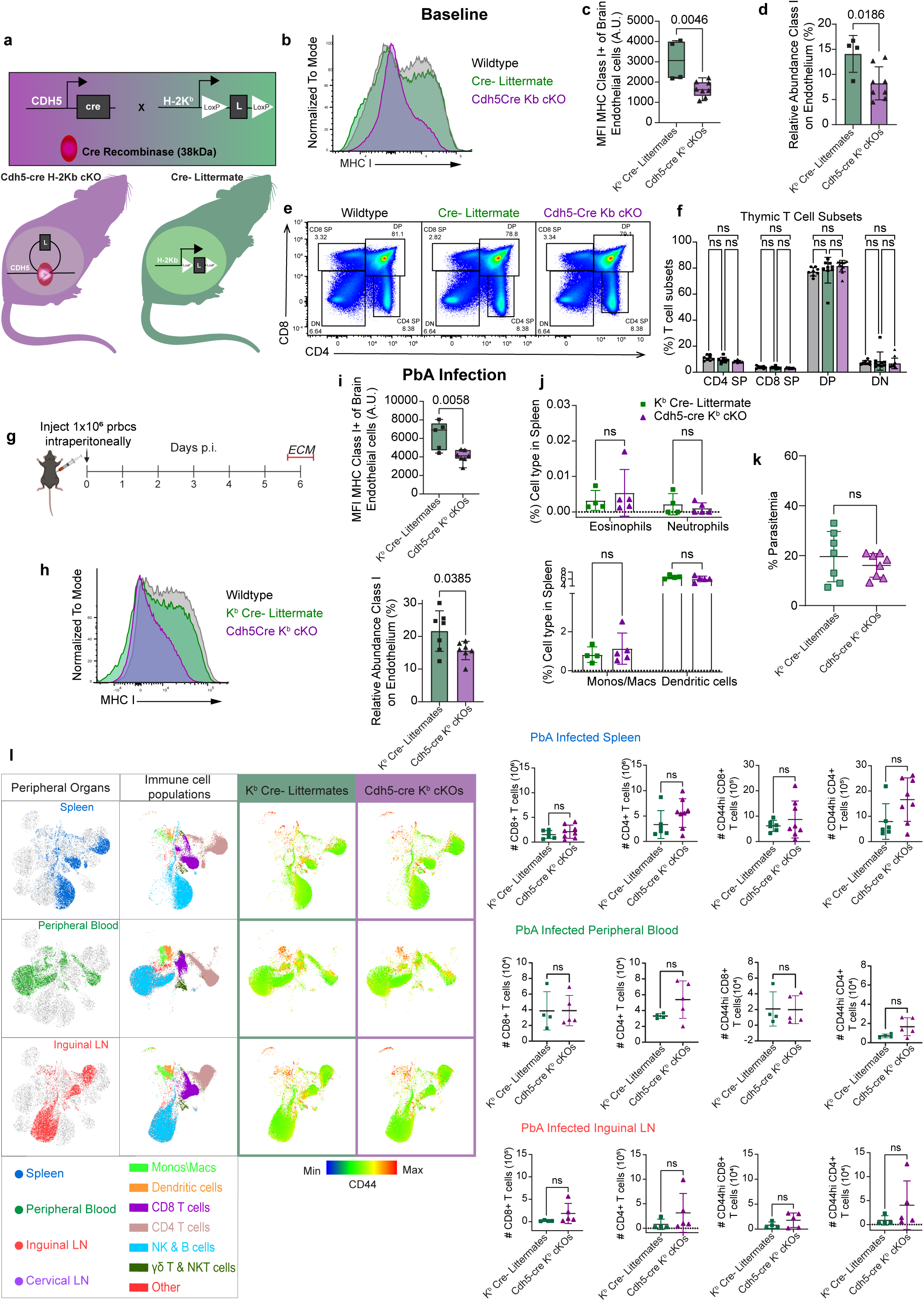
Normal peripheral infection and immunity in mice with K^b^ class I molecule conditionally ablated on vasculature. a) Schematic depicted Cre-LoxP strategy. Cdh5-Cre mice were crossed to transgenic H-2K^b^ LoxP mice. b) Representative MFI plot comparing C57BL/6, Cdh5-Cre K^b^ cKO, and Cre-Littermate expression of MHC I on brain endothelium in naïve mice. c) Quantification of MHC I MFIs on brain endothelial cells using arbitrary units (A.U.) in naïve mice. d) Percentage of brain endothelial cells expressing high levels of MHC Class I molecules in naïve mice. e) Representative flow plot of C57BL/6, Cdh5-Cre K^b^ cKO, and Cre-Littermate T cell subsets in thymus. f) Percentage of thymic T cell subsets. g) Schematic representing timeline of infection for development of ECM. h) Representative MFI plot comparing C57BL/6, Cdh5- Cre K^b^ cKO, and Cre-Littermate expression of MHC I on brain endothelium in PbA infected mice. i) Quantification of MHC I MFIs on brain endothelial cells using arbitrary units (A.U.) in PbA infected mice. j) Percentage of brain endothelial cells expressing high levels of MHC Class I molecules in PbA infected mice. k) Percentage of granulocytes in spleens of infected mice. l) Percentage of antigen-presenting cells in spleens of infected mice. m) Percent parasitemia quantified by imaging Giemsa-stained thin blood smears in PbA infected mice. n) Representative UMAPs depicting peripheral organs and immune cell populations and respective heats maps with CD44 activation marker expression. o) The scatter plots show quantified absolute cell counts of CD8 T cells, CD4 T cells and CD44hi CD8 T cells and CD4 T cells in the respective organs depicted in n). Data were analyzed with unpaired t test or 2way ANOVA with multiple comparisons. Scatterplots and bar plots present the mean ± SD. All p-values are shown or ns for non-significant p-value *(p > 0.05)*.

### Endothelial cell specific D^b^ conditional knockout mice develop normal peripheral *Plasmodium* infection and immune responses

We crossed (class I deficient background, K^b^−/−, D^b^−/−) D^b^ LoxP mice with (class I deficient background, K^b^−/−, D^b^−/−) Cdh5-Cre mice (Fig. 3a). The resulting animals are defined as either D^b^ Cre- littermate (with ubiquitous D^b^ expression), or Cdh5-Cre D^b^ cKO (with endothelial-specific ablation of D^b^). Endothelial cell expression of D^b^ at baseline in Cdh5-Cre D^b^ cKO mice was comparable to Cre- littermate controls. Percentage of class I expression on endothelium was reduced but did not reach statistical significance in uninfected Cdh5-Cre D^b^ cKO mice (Fig. 3b-d). T cell development was intact with no changes in thymic T cell subset frequencies, mTEC and cTEC frequencies, or in thymic weights between groups (Fig. 3e-f and Supplementary Fig. 4a). There were no differences between males and females in development of thymic T cell subsets, and no differences between groups in CD8 T cell V beta usage in spleen, or in splenic weights (Supplementary Fig. 4b-d). To determine the effect of D^b^ class I molecule ablation on MBMECs during ECM, adult Cdh5-Cre D^b^ cKO (∼3-month-old) mice were I.P. infected with PbA and tissues were harvested at onset of severe neurological symptoms, approximately 6-8 dpi (Fig. 3g). Importantly, during infection D^b^ expression was significantly reduced on brain endothelium (Fig. 3h-j). Spleens of these animals had no changes to granulocyte or APC subsets during infection (Fig. 3k-l) and mice established full blood stage infection in the periphery (Fig. 3m). Assessment of immune cell populations in peripheral tissues at baseline and during infection revealed no changes in CD45+ immune cell number between D^b^ Cre- littermates and Cdh5-Cre D^b^ cKO in spleen, peripheral blood or inguinal LN (Supplementary Fig. 4e-f). CD4 and CD8 T cell numbers and activation state remained unchanged between groups in all peripheral organs at baseline (Supplementary Fig. 4f). However, during infection, there was reduction in number of activated CD8 T cells in inguinal LN in the Cdh5-Cre D^b^ cKO animals (Fig. 3n-p). These results demonstrate that our Cdh5-Cre D^b^ cKO mouse model has sufficient deletion of D^b^ on vasculature during PbA infection and a fully functional peripheral immune response. These results enabled us to proceed with interrogation of potentially brain-specific immunomodulatory functions of D^b^ Ag-presentation by brain endothelial cells in ECM.

**Fig 3.**
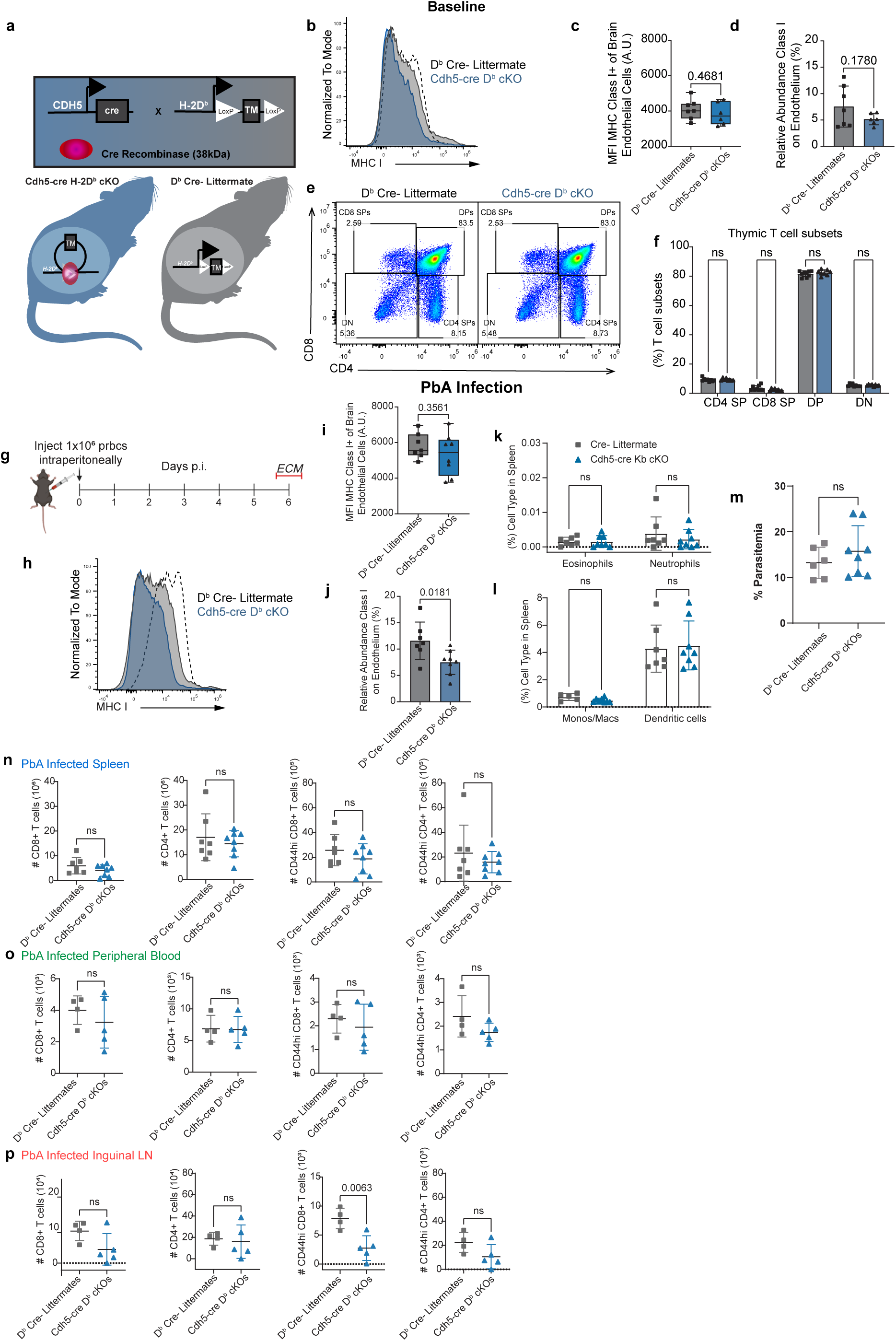
Normal peripheral infection and immunity in mice with D^b^ class I molecule conditionally ablated on vasculature. a) Schematic depicted Cre-LoxP strategy. Cdh5-Cre mice were crossed to transgenic H-2D^b^ LoxP mice. b) Representative MFI plot comparing C57BL/6, Cdh5-Cre D^b^ cKO, and Cre-Littermate expression of MHC I on brain endothelium in naïve mice. c) Quantification of MHC I MFIs on brain endothelial cells using arbitrary units (A.U.) in naïve mice. d) Percentage of brain endothelial cells expressing high levels of MHC Class I molecules in naïve mice. e) Representative flow plot of C57BL/6, Cdh5-Cre D^b^ cKO, and Cre-Littermate T cell subsets in thymus. f) Percentage of thymic T cell subsets. g) Schematic representing timeline of infection for development of ECM. h) Representative MFI plot comparing C57BL/6, Cdh5- Cre D^b^ cKO, and Cre-Littermate expression of MHC I on brain endothelium in PbA infected mice. i) Quantification of MHC I MFIs on brain endothelial cells using arbitrary units (A.U.) in PbA infected mice. j) Percentage of brain endothelial cells expressing high levels of MHC Class I molecules in PbA infected mice. k) Percentage of granulocytes in spleens. l) Percentage of antigen-presenting cells in spleens. m) Percent parasitemia quantified by imaging Giemsa-stained thin blood smears in PbA infected red blood cells. n) Scatter plots show quantified absolute cell counts of CD8 T cells, CD4 T cells and CD44hi CD8 T cells and CD4 T cells in spleen. o) Scatter plots show quantified absolute cell counts of CD8 T cells, CD4 T cells and CD44hi CD8 T cells and CD4 T cells in peripheral blood. p) Scatter plots show quantified absolute cell counts of CD8 T cells, CD4 T cells and CD44hi CD8 T cells and CD4 T cells in inguinal LN. Data were analyzed with unpaired t test or 2way ANOVA with multiple comparisons. Scatterplots and bar plots present the mean ± SD. All p-values are shown or ns for non-significant p-value *(p > 0.05)*.

### T cell dynamics at the barrier are modulated by the presence of distinct class I molecules on endothelium

To assess whether brain vasculature was inherently changed by loss of endothelial class I in cKO mice, we performed maximum intensity projections (MIPs) using MRI angiography to illuminate and assess the blood vessels. Cdh5-Cre K^b^ cKO and Cdh5-Cre D^b^ cKO were compared to their respective Cre- littermate controls. At baseline, we observed no differences in 3D volume of overall brain vasculature or voxel units (Supplementary Fig. 5a-b and 5c-d). We therefore proceeded to assess the effects of discrete MHC class I molecules on T cell dynamics in brain vasculature. To address this, we employed acute cranial window placement and *in vivo* two-photon imaging after IV injections of anti-Thy1.2-Alexa Fluor 488 to label T cells in vasculature and Texas Red-Dextran to label vessels. 6-7 days following PbA infection, mice underwent live imaging and T cell interactions at the brain vasculature were tracked over time (Fig. 4a). Cre- Littermates with K^b^ expression on brain vasculature presented with higher numbers of T cells per field of view (FOV) with slowed velocities, indicating interactions with K^b^ on endothelium (Fig. 4b and 4c). In contrast, Cdh5-Cre K^b^ cKO mice, which lack K^b^ on endothelium, had lower numbers of T cells per FOV, with higher velocity indicating less interactions with endothelium (Fig. 4b and 4c). Interestingly, there were no significant differences between the frequency that T cells contacted the vessels between genotypes. However, arrest on vessels was significantly increased when K^b^ was expressed on the brain vasculature (Fig. 4d-4e and Supplementary Videos 1-2). To investigate these T cell interactions further, we examined the activation state and adhesion molecule expression of MBMECs. During late-stage PbA infection, endothelial cells upregulate adhesion molecules responsible for initiating T cell extravasation^20^. P-selectin, which facilitates activated T cell rolling, did not differ in expression between Cdh5-Cre K^b^ cKO mice and Cre- littermates (Fig. 4f). ICAM-1 and VCAM-1 expression, involved in arrest and adhesion, also showed no significant differences during infection (Fig. 4g-h). When endothelial cells are activated, they upregulate molecules involved in Ag-presentation^17^. Surprisingly, when K^b^ class I molecule expression was ablated, MHC Class II molecules were upregulated on brain endothelium during infection (Fig. 4i).

**Fig 4.**
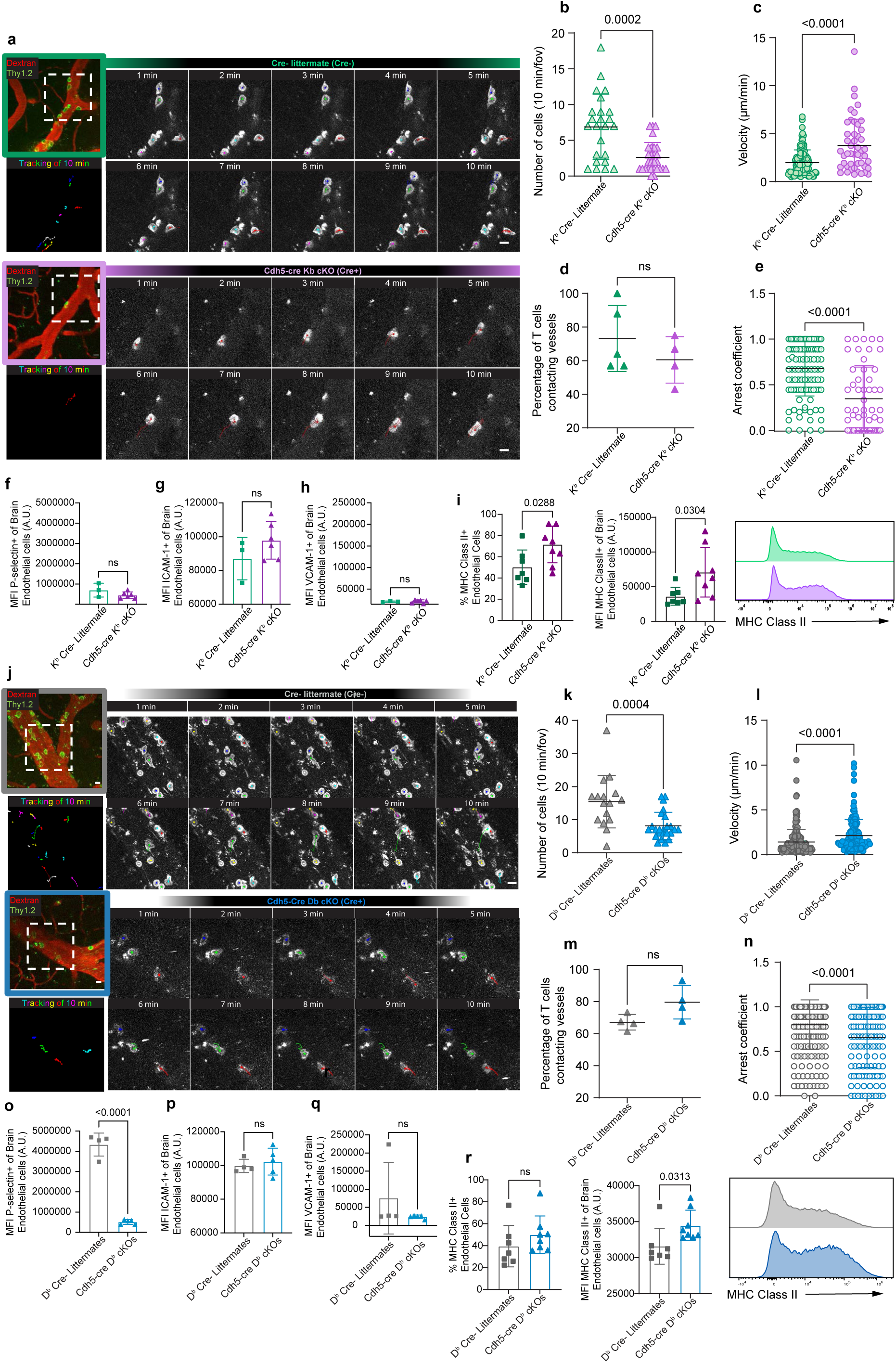
T cell dynamics at brain vasculature are dependent on MHC class I expression during infection. a) Representative images of T cells in brain vasculature with blood vessels labeled with I.V. Dextran-Texas ∼5 min. prior to imaging and T cells I.V. labeled with anti-Thy1.2-Alexa Fluor 488 ∼30 min. Inset is representative of T cell tracking over time through acute cranial window on days 6-7 post PbA infection. T cells (green) were visualized interacting with blood vessels (red) over periods of 10-30 min. Upper row represents K^b^ Cre- littermates and bottom row represents Cdh5-cre K^b^ cKO mice. Scale bar, 10 µM. b) Scatter plot represents numbers of T cells visualized over 10 min time periods/FOV. c) T cell velocity was measured using the Manual Tracking plugin in FIJI. Data are representative of µm/min over periods of 10 min per field and per cell. d) Percentage of T cells contacting vessels was calculated by obtaining the percent of all visualized T cells that contacted vasculature over many fields of view. Each data point is representative of the overall average percentage/mouse. e) The arrest coefficient is the percentage of each cell track that a cell is immobile (instantaneous velocity <2 µm/min). Each dot represents one cell and its percentage of contacts over time. (n = 5-7 mice per group). See corresponding S1-S5 movies. High-dimensional flow cytometry was performed to assess levels of protein expression on brain endothelium day 6 of PbA infection. f) Scatter plot represents MFI values of P-selectin on brain endothelial cells. g) MFI of ICAM-1on brain endothelial cells h) MFI of VCAM-1 on brain endothelial cells. i) Percentage and MFI quantification of MHC Class II molecules on brain endothelial cells with representative histogram. j) Representative images of T cells in brain vasculature labeled as described above. Inset is representative of T cell tracking over time through acute cranial window on days 6-7 post PbA infection. T cells (green) were visualized interacting with blood vessels (red) over periods of 10-30 min. Upper row represents D^b^ Cre- littermates and bottom row represents Cdh5-cre D^b^ cKO mice. Scale bar, 10 µM. k) Scatter plot represents numbers of T cells visualized over 10 min time periods/FOV. l) T cell velocity was measured using the Manual Tracking plugin in FIJI. Data are representative of µm/min over periods of 10 min per field and per cell. m) Percentage of T cells contacting vessels was calculated by obtaining the percent of all visualized T cells that contacted vasculature over many fields of view. Each data point is representative of the overall average percentage/mouse. n) The arrest coefficient is the percentage of each cell track that a cell is immobile (instantaneous velocity <2 µm/min). Each dot represents one cell and its percentage of contacts over time. (n = 5 mice per group). See corresponding S1-S5 movies. High-dimensional flow cytometry was performed to assess levels of protein expression on brain endothelium day 6 of PbA infection. o) Scatter plot represents MFI values of P-selectin on brain endothelial cells. p) MFI of ICAM-1on brain endothelial cells q) MFI of VCAM-1 on brain endothelial cells. r) Percentage and MFI quantification of MHC Class II molecules on brain endothelial cells with representative histogram. Data were analyzed with unpaired t test. Scatterplots and bar plots present the mean ± SD. All p-values are shown or ns for non-significant p-value *(p > 0.05)*.

T cell dynamics in Cre- D^b^ expressing and Cdh5-Cre D^b^ cKO mice were similar to those of the previous model. 6-7 dpi with PbA T cell interactions with vasculature were tracked over time during intravital imaging (Fig. 4j). Cre- Littermates with D^b^ present on vasculature showed high numbers of T cells/FOV. In Cdh5-Cre D^b^ cKO mice, T cells were greatly reduced per FOV, indicating fewer T cells slowed down to interact with endothelium when D^b^ molecule is ablated on vasculature (Fig. 4k). Meanwhile, D^b^ expressing Cre- littermate T cells had slower velocities, indicating more interactions with D^b^ on endothelium. When endothelial D^b^ is conditionally deleted in Cdh5-Cre D^b^ cKO mice, T cells that interact with vessels had significantly increased velocities, with reduced slowing and rolling on vasculature (Fig. 4l). Again, there were no significant differences between the percentage of T cells contacting vessel walls between groups. However, the frequency of T cell arrests was significantly increased when D^b^ was expressed by brain vasculature (Fig. 4m and 4n). (Supplementary Videos 2-3). P-selectin was greatly increased when D^b^ was present on brain endothelium during infection, indicating a higher probability of activated T cell rolling (Fig. 4o). Interestingly, cells/FOV in the D^b^ expressing Cre- littermates, had increased cells/FOV compared to K^b^ Cre- littermates (Fig. 4b and 4k). There were no significant changes in ICAM-1 or VCAM-1 expression on endothelium when D^b^ was conditionally deleted (Fig. 4p-q). As was seen with K^b^ cKO mice, deletion of D^b^ class I molecule on endothelial cells resulted in higher expression of MHC Class II molecules on endothelium in comparison to Cre- littermates (Fig.4i and 4r). Together, this data demonstrates that H-2K^b^ and H-2D^b^ class I molecules are critical for optimal T cell adhesion and arrest to brain endothelium during ECM.

### CD8 T cell activation and infiltration into brain parenchyma is differentially modulated by discrete class I molecules during infection

We next addressed if the changes observed in T cell dynamics at the brain vasculature during PbA infection corresponded with T cell infiltration into brain parenchyma. First, we wanted to confirm that cervical lymph node (cLN) priming of brain draining antigens was intact in our mouse models. To test this, we employed Theiler’s murine encephalomyelitis virus (TMEV) and TMEV-OVA. These neurotropic viruses are intracranially administered and provide a highly reproduceable model of CNS antigen drainage resulting in cLN priming, CD8 T cell infiltration, and viral clearance^21, 22, 23^. TMEV immunodominant peptide VP2_121-130_ is presented in the H-2D^b^ MHC class I molecule whereas TMEV-OVA model antigen OVA_257-264_ is presented by H-2K^b^ ^23^. On day 7 post infection mice were perfused and cLNs were analyzed via spectral flow cytometry (Supplementary Fig. 6a). In mice infected with either TMEV or TMEV-OVA, tetramer staining was unchanged between Cre- and Cre+ mice indicating that cLN priming in these mice is intact (Supplementary Fig. 6b-c). We now evaluated immune cell infiltration into the brain in the ECM model, in which the *Plasmodium* infection arises in the periphery prior to neuroinflammation in the brain (Fig. 5a). At moribund, 6-8 d.p.i., mice were euthanized and perfused. Mouse brains were processed into single cell suspensions and analyzed via spectral flow cytometry. Overall immune populations in the brains during infection can be visualized in the UMAP projection (Fig. 5b). Data encompassing CNS infiltration of immune cells were interrogated. Absolute counts of total CD45+ immune cells in the brain remain unchanged when K^b^ class I molecules are present on endothelium versus endothelial cell class I molecule conditional knockout mice (Supplementary Fig. 6d). Numbers of CD4 T cells in the brain also remain unchanged regardless of K^b^ expression by endothelial cells (Fig. 5e). However, when K^b^ expression is lost on endothelium, CD8 T cell infiltration of the brain is greatly diminished (Fig. 5f). Since adhesion molecule expression on endothelium was previously shown to be unchanged, we also examined CD8 T cells for chemokine receptors involved in trafficking to the CNS and adhesion to the endothelium. Levels of CXCR3, a receptor shown to be necessary for recruitment of T cells into the brain to establish ECM^24, 25^, was not impacted by ablation of K^b^ on vasculature (Supplementary Fig. 6f-g). CCR7, a chemokine receptor known to be upregulated on T cells trafficking to the brain during neuroinflammation, was also evaluated^26^. During ECM the olfactory region highly upregulates CCL21, and blockade of CCR7 decreases T cell activation, recruitment and survival^27^. In Cre- littermates with K^b^, we saw no difference in CD8 T cell expression of CCR7 from mice lacking K^b^ on vasculature (Supplementary Fig. 6h-i). We next evaluated effector status of CD4 and CD8 T cells in the brain. Interestingly, CD8 T cells that infiltrated the brain during infection produce significantly less IFN-y when K^b^ expression was ablated on vasculature (Fig. 5g). Surprisingly, although brain infiltrating CD4 T cell numbers did not change, these lymphocytes increasingly assumed a Th1 phenotype with increased expression of IFN-y when K^b^ was ablated on endothelium (Fig. 5h). In a parallel set of experiments, we investigated whether antigen recognition plays a role in infiltration and localization of IFN-y producing CD8 T cells. We infected mice and intravenously (IV) injected Alexa Fluor-488 conjugated anti-Thy1.2 antibody 3 minutes before euthanasia on day 6 of infection to label cells that have recently infiltrated brain parenchyma from the bloodstream. It is important to note that K^b^ expressing Cre- mice are not yet moribund on day 6 of infection. We also evaluated surface expression of CD11a as a surrogate marker for cognate antigen-experienced CD8 T cells as previously described ^28^. At 6 d.p.i. we show that numbers of Ag-experienced CD8 T cells were not different between K^b^ expressing Cre- and Cdh5-Cre K^b^ cKO mice in meninges or cervical LNs of PbA infected mice. However, numbers of brain infiltrated Ag-specific CD8 T cells were greatly reduced when K^b^ was lost on vasculature (Fig.5i). K^b^ expressing Cre- mice presented with a significant increase of CD11a+ CD8 T cells only in the IV- group, indicating that the majority had infiltrated the brain prior to IV labeling. In Cdh5-Cre K^b^ cKO mice lacking K^b^ on vessels, numbers of both IV- and IV+ CD8 T cells were greatly reduced, demonstrating that recognition of peptides presented by K^b^ is required for optimal CD8 T cell infiltration (Fig. 5j). It is known that IFN-y producing CD8 T cells have enhanced Ag-specific contacts, cytolytic killing and kinetics through paracrine and autocrine IFN-y signaling^29^. We therefore employed IV labeling to determine the localization of IFN-y producing CD8 T cells 6 d.p.i with PbA. We noted a slight trend towards increased frequency of IFN-y producing CD8 T cells in the recently infiltrated IV+ population in both Cre- and Cre+ animals. However, this difference did not reach significance (Fig5k).

**Fig 5.**
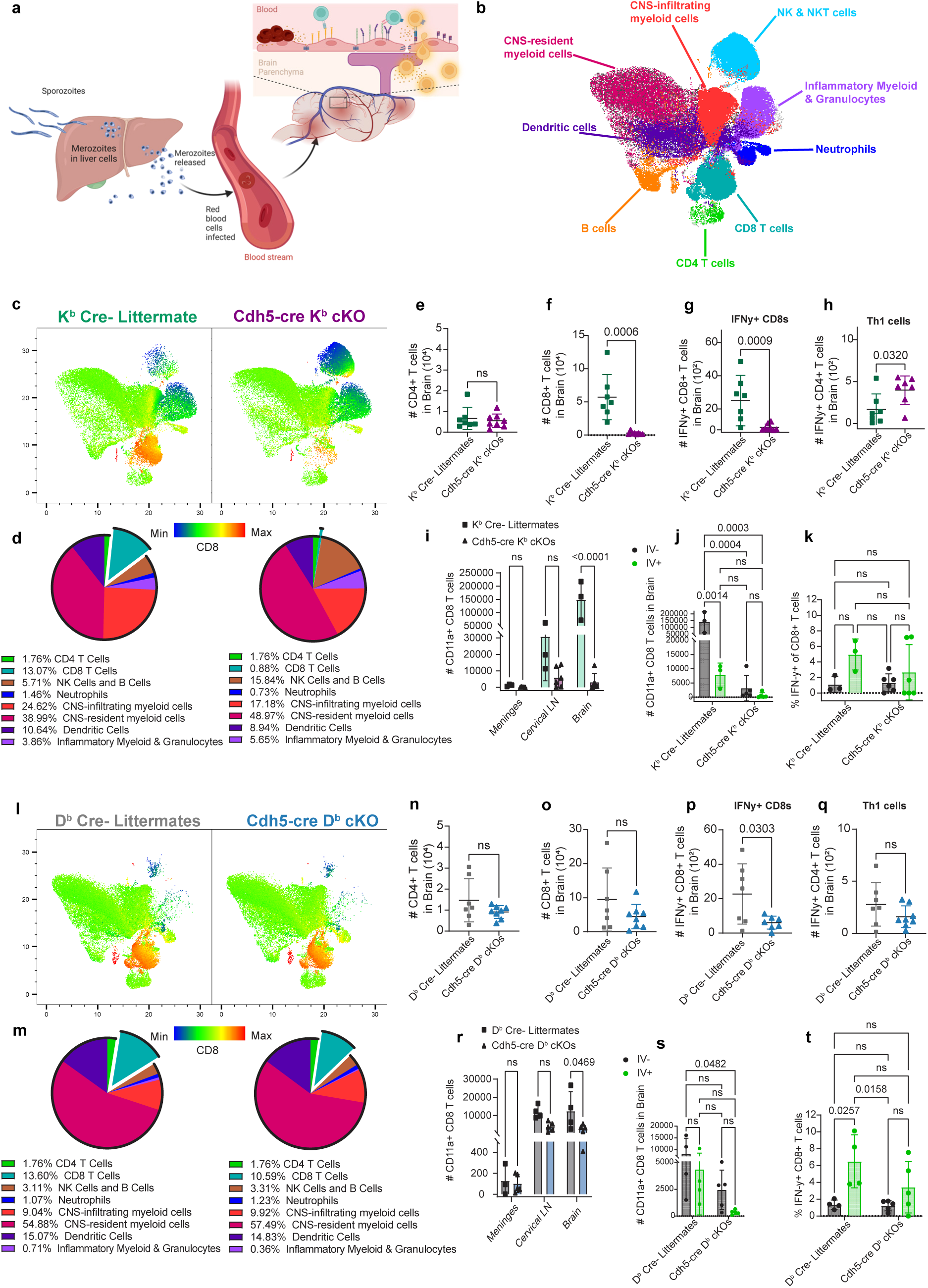
Modulation of individual class I molecules on brain endothelium results in brain-specific CD8 T cell infiltration changes. a) BioRender schematic of events in the periphery leading up to cerebral malaria. b) Representative UMAP key, colorized to highlight the different cell subsets in the brain. c) Representative UMAPs of K^b^ Cre- Littermate and Cdh5-cre K^b^ cKO brains, colorized with heat map of CD8 expression. d) Pie charts with corresponding frequencies of cell subset in K^b^ Cre- Littermate and Cdh5-cre K^b^ cKO brains (of CD45+) e) Absolute counts of CD4 T cells in the brains of infected K^b^ Cre- Littermate and Cdh5-cre K^b^ cKO brains. f) Absolute counts of CD8 T cells in the brains of infected K^b^ Cre- Littermate and Cdh5-cre K^b^ cKO brains. g) Absolute counts of IFN-y+ CD8 T cells in the brains of infected K^b^ Cre- Littermates and Cdh5-cre K^b^ cKOs. h) Absolute counts of IFN-y+ CD4 T (Th1) cells in the brains of infected K^b^ Cre- Littermates and Cdh5-cre K^b^ cKOs. i) Absolute counts of CD11a+ (antigen-experienced) CD8 T cells in the meninges, cLN and brain of infected K^b^ Cre- Littermates and Cdh5-cre K^b^ cKOs. Mice were injected with Alexa Fluor 488-anti-Thy1.2 antibody ∼3 min. before euthanasia. j) Quantified absolute counts of either IV+ or IV- antigen-experienced CD8 T cells in the brains of infected K^b^ Cre- Littermates and Cdh5-cre K^b^ cKOs. k) Frequency of IFN-y+ CD8 T cells either labeled with IV antibody or negative for IV labeling. l) Representative UMAPs of D^b^ Cre- Littermate and Cdh5-cre D^b^ cKO brains, colorized with heat map of CD8 expression. m) Pie charts with corresponding frequencies of cell subset in D^b^ Cre- Littermate and Cdh5-cre D^b^ cKO brains (of CD45+) n) Absolute counts of CD4 T cells in the brains of infected D^b^ Cre- Littermate and Cdh5-cre D^b^ cKO brains. o) Absolute counts of CD8 T cells in the brains of infected D^b^ Cre- Littermate and Cdh5-cre D^b^ cKO brains. p) Absolute counts of IFN-y+ CD8 T cells in the brains of infected D^b^ Cre- Littermates and Cdh5-cre D^b^ cKOs. q) Absolute counts of IFN-y+ CD4 T (Thl) cells in the brains of infected D^b^ Cre- Littermates and Cdh5-cre D^b^ cKOs. r) Absolute counts of CD11a+ (antigen-experienced) CD8 T cells in the meninges, cLN and brain of infected D^b^ Cre- Littermates and Cdh5-cre D^b^ cKOs. Mice were injected with Alexa Fluor 488-anti-Thy1.2 antibody ∼3 min. before euthanasia. s) Quantified absolute counts of either IV+ or IV- antigen-experienced CD8 T cells in the brains of infected D^b^ Cre- Littermates and Cdh5-cre D^b^ cKOs. t) Frequency of IFN-y+ CD8 T cells either labeled with IV antibody or negative for IV labeling. Data were analyzed with unpaired t test or 2way ANOVA with multiple comparisons. Scatterplots and bar plots present the mean ± SD. All p-values are shown or ns for non-significant p-value *(p > 0.05)*.

We next assessed the immune profile of the brains of D^b^ expressing mice in ECM. Tissues were harvested at moribund, days 6-8 of infection. We observed no difference in the overall numbers of CD45+ immune cells in the brain between D^b^ expressing Cre- and Cdh5-Cre D^b^ cKO mice (Supplementary Fig. 6e). Surprisingly, we observed no change in CD8 T cell infiltration between the groups, and numbers of CD4 T cells in the brain were also unchanged (Fig. 5l-o). Expression of trafficking and adhesion molecules on CD8 T cells were assessed and levels of CXCR3 and CCR7 were not significantly different between D^b^ Cre- littermates and Cdh5-Cre D^b^ cKO (Supplementary Fig. 6j-m). Although overall brain infiltration of CD8 T cells was unchanged, these lymphocytes presented with reduced IFN-y production when D^b^ was ablated on vasculature (Fig. 5p). In contrast, no change in CD4 T cell IFN-y production was observed (Fig. 5q). When assessing the role of Ag-recognition in infiltration, we observed 6 d.p.i. that numbers of CD11a+ Ag-experienced CD8 T cells were not different between Cre- and Cdh5-Cre D^b^ cKO mice in meninges or cervical LNs. However, numbers of brain infiltrating CD11a+ Ag-experienced CD8 T cells were significantly reduced when D^b^ was lost on vasculature (Fig. 5r). We observed that numbers of previously infiltrated, IV- Ag-experienced CD8 T cells were much lower in D^b^ Cre- mice than in K^b^ Cre- mice, although the IV+ populations between Cre-s were more comparable on day 6 of infection (Fig. 5j and 5s). This suggests that Ag-specific CD8 T cell infiltration may begin at a later timepoint and escalate more quickly with D^b^ on endothelium in comparison to K^b^. Alternatively, we may not see a difference between IV- and IV+ if vascular leakage is present as this could result in increased antibody staining in the parenchyma (Fig. 5s). The previously infiltrated, Ag-experienced CD8 T cells in the D^b^ Cre- littermates were significantly increased only when compared to the IV+ population in the D^b^ cKO mice (Fig. 5s).

It is important to highlight that total numbers of brain infiltrated CD8 T cells are similar between Cre- littermates (with either K^b^ or D^b^) at endpoint (Fig. 5f and 5o). However, Ag-experienced CD11a+ CD8 T cell numbers were much higher in K^b^ expressing than in D^b^ expressing Cre- controls (Fig. 5i and 5r). As per localization of IFN-y producing CD8 T cells, D^b^ expressing Cre- animals had significantly increased frequency of CD8 T cells producing IFN-y in the IV labeled population on day 6 of infection, suggesting these cells have increased effector function when recently infiltrated from vasculature (Fig. 5t). Meanwhile, Cdh5-Cre D^b^ cKO animals had no change in frequency of IFN-y producing CD8 T cells between IV+ and IV- groups (Fig. 5t).

To further interrogate infiltration, we compared rates of entry of Ag-experienced (CD11a+) and Ag-inexperienced (CD11a-) CD8 T cells on day 6 of infection. We observed that CD11a+ Ag-experienced CD8 T cells in mice with K^b^ or D^b^ expression had faster infiltration than mice with conditional deletion of class I molecules on endothelial cells, though this did not reach significance in D^b^ cKO mice (Supplementary Fig. 6n-o). CD8 T cells that were CD11a- (Ag-inexperienced) in K^b^ expressing mice had low rates of infiltration and there was not a significance difference in infiltration between K^b^ Cre- and Cdh5-Cre K^b^ cKO mice (Supplementary Fig. 6p). These data suggest that K^b^ class I molecule is important for Ag-specific CD8 T cell brain infiltration in ECM. Although no differences were seen in CD11a- Ag- inexperienced CD8 T cell rates of entry in between Cre- and D^b^ cKO mice, D^b^ expressing Cre- mice rate of CD11a- CD8 T cell entry was double that of K^b^ expressing Cre- animals (Supplementary Fig. 6p and q). Importantly, the same number of activated IFN-y producing CD8 T cells end up in the brain of K^b^ and D^b^ Cre- littermates at endpoint despite numbers of CD11a+ Ag-experienced CD8 T cells in D^b^ expressing mice being much lower on day 6 (Fig. 5g and p). This demonstrates that D^b^ class I molecule on endothelium is required to initiate IFN-y production by CD8 T cells to the same extent as K^b^, and that each class I molecule on endothelium differentially modulates the timeline of entry of Ag-specific CD8 T cells.

It is known that most neuropathology and vascular leakage during CM and ECM localize to three distinct regions of the brain; olfactory, hippocampus and brainstem^16^. We therefore wanted to assess whether infiltrating CD8 T cells enter brain parenchyma in these regions. Using 2-photon and confocal microscopy, we imaged cell dynamics and performed immunofluorescence staining on brain tissues at late stages of infection. At 6-7 dpi we observed CD8 T cells extravasating via post-capillary venules and entering regions of brain parenchyma in K^b^ expressing Cre- mice (Supplementary Video 5). In K^b^ expressing Cre- littermates, we observed parenchymal CD8 T cells in olfactory, brainstem, and hippocampal regions. However, when K^b^ is deleted on vasculature, CD8 T cell infiltration into these regions is greatly decreased (Supplementary Fig. 6r). When imaging D^b^ expressing Cre- littermates, we saw CD8 T cell infiltration into brain parenchyma (Supplementary Video 6). We also confirmed that when D^b^ expression is ablated on vasculature, CD8 T cells are still present in brain parenchyma in these three regions (Supplementary Fig. 6s). These data support the findings of our spectral flow cytometry data and demonstrate that CD8 T cells are infiltrated into regions of the brain affected by ECM, except specifically when K^b^ is deleted on brain vasculature.

### Blood-brain barrier disruption is dependent on MHC Class I molecule expression by brain endothelium

Next, we assessed the contribution of discrete class I molecules on modulating BBB disruption during ECM. At 6-8 dpi with PbA, mice were administered gadolinium I.P. and evaluated for vascular permeability in the brain using T1-weighted gadolinium enhanced MRI. In Cre- mice with K^b^ class I molecule expression intact, we see significant enhancement in regions of olfactory bulb, brainstem and hippocampal regions periventricular to the lateral ventricles (Fig 6a). All areas of leakage were assessed utilizing 3D volumetric analysis with Analyze Software as previously described^16^. Brain vascular integrity was retained, and gadolinium leakage was not observed when K^b^ expression was ablated on vasculature (Fig. 6a-c). As a secondary method of vascular leakage assessment, we IV injected ∼66 kDa FITC-albumin, one hour prior to tissue harvest. FITC-albumin mean fluorescence intensity was quantified in our three brain regions of interest. Uninfected and infected wildtype C57BL/6 mice were included as visual reference controls (Fig. 6d). Cre- littermates with K^b^ expression had significant FITC-albumin leakage in regions of olfactory bulb and brainstem. In contrast, Cdh5- Cre K^b^ cKO animals lacking K^b^ on vasculature had negligible BBB disruption, indistinguishable from uninfected controls (Fig. 6d). FITC-albumin leakage is quantified in Fig. 6e. PbA infected D^b^ expressing Cre- Littermates presented with even more pronounced gadolinium-enhancement in regions of interest than K^b^ expressing Cre- mice as measured by small animal MRI (Fig. 6a-c and 6f-h). Cdh5-Cre D^b^ cKO mice however, presented with an intact blood brain barrier, with no notable pathology when D^b^ is lost on vasculature during infection (Fig. 6f-h). In a secondary analysis utilizing FITC-albumin leakage, we observed increased FITC-albumin mean fluorescence intensity in D^b^ expressing Cre- olfactory bulb and brainstem (Fig. 6i-j). In Cdh5- Cre D^b^ cKO mice, brain vascular integrity was retained, and no FITC-albumin leakage was observed (Fig. 6i-j). Tight junction proteins are known to be downregulated in regions of BBB disruption^16^. In human endothelial cells exposed to *Plasmodium falciparum* prbcs *ex vivo*, we observed negative enrichment scores consistent with downregulation of tight junction proteins (Supplementary Fig. 7a). We next performed Claudin-5 staining in our K^b^ and D^b^ models, interrogating the regions of vascular leakage in the brains, where T cells were also observed to infiltrate (Supplementary Fig. 8r-s). We saw that the vessels in regions of leakage and CD8 T cell infiltration in both K^b^ and D^b^ Cre- littermates had a more sparse and punctate staining for Claudin-5, indicating tight junction protein alteration (Supplementary Fig. 7b-c). However, K^b^ and D^b^ cKO animals lacking class I molecules on vasculature had intact Claudin-5 expression along vessels, despite the CD8 T cell infiltration seen in the Cdh5-Cre D^b^ cKO mice (Supplementary Fig. 7b-c). Overall, these results indicate that vascular leakage and disruption of BBB tight junctions occurs in mice via class I molecule restricted antigen presentation by brain endothelium during *Plasmodium* infection.

**Fig 6.**
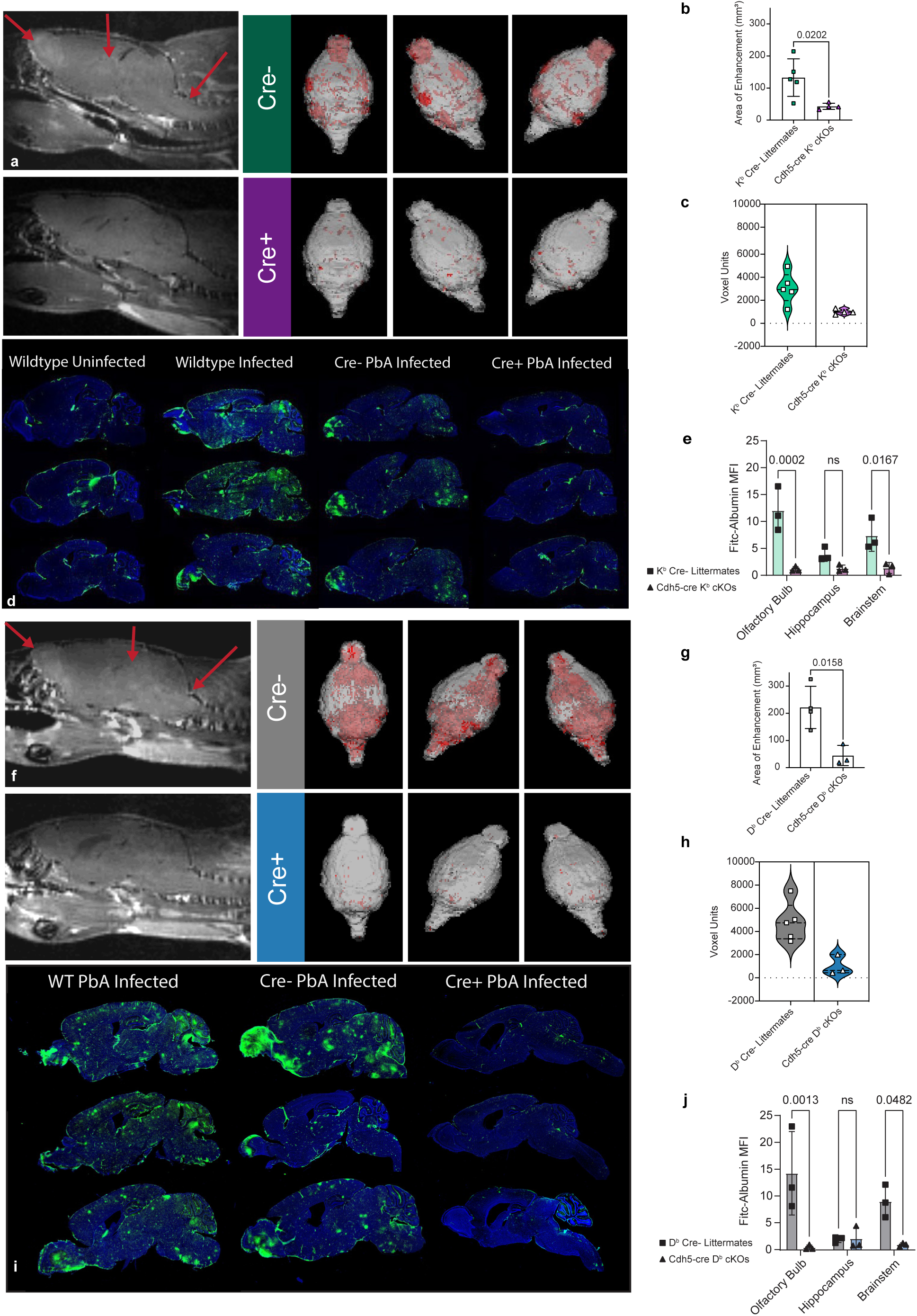
Vascular leakage during PbA infection is dependent on class I molecule expression on endothelium. a) Top – Representative T1-weighted gadolinium enhanced MRI (sagittal slice) from K^b^ Cre- littermate day 6-8 PbA infection showing vascular leakage in regions of enhancement. Bottom - Representative T1-weighted gadolinium enhanced MRI (sagittal slice) from Cdh5-cre K^b^ cKO mice day 6-8 PbA infection with no vascular leakage or enhancement noted. Top right – 3D volumetric modeling of whole brain with vascular leakage highlighted in red. Top left – 3D volumetric modeling of whole brain with no apparent vascular leakage noted. b) Area measurements of regions of enhancement. Enhancement was quantified for each individual MRI slice and combined into a cumulative measurement per mouse. 2 blinded individuals performed measurements. Each data point represents the average of the scores for each mouse/scan. c) 3D pixelated volumetric measurement were obtained from all slices and averaged from 2 blinded reviewers as mentioned in b. d) FITC-albumin leakage assay results. Mice were IV injected with 66kDa FITC-conjugated albumin 1 hour prior to euthanasia to visualize regions of blood-brain barrier disruption and vascular leakage. e) Quantification of FITC-albumin mean fluorescence intensity per ROI. f) Top – Representative T1-weighted gadolinium enhanced MRI (sagittal slice) from D^b^ Cre- littermate day 6-8 PbA infection showing vascular leakage in regions of enhancement. Bottom - Representative T1-weighted gadolinium enhanced MRI (sagittal slice) from Cdh5-cre D^b^ cKO mice day 6-8 PbA infection with no vascular leakage or enhancement noted. Top right – 3D volumetric modeling of whole brain with vascular leakage highlighted in red. Top left – 3D volumetric modeling of whole brain with no apparent vascular leakage noted. g) Area measurements of regions of enhancement. Enhancement was quantified for each individual MRI slice and combined into a cumulative measurement per mouse. 2 blinded individuals performed measurements. Each data point represents the average of the scores for each mouse/scan. h) 3D pixelated volumetric measurement were obtained from all slices and averaged from 2 blinded reviewers as mentioned in g. i) FITC-albumin leakage assay results. Mice were IV injected with 66kDa FITC-conjugated albumin 1 hour prior to euthanasia to visualize regions of blood-brain barrier disruption and vascular leakage. j) Quantification of FITC-albumin mean fluorescence intensity per ROI. Data were analyzed with unpaired t test or 2way ANOVA with multiple comparisons. Violin plots and bar plots present the mean ± SD. All p-values are shown or ns for non-significant p-value *(p > 0.05)*.

### PbA infected parabionts sharing peripheral circulation demonstrate critical importance of endothelial D^b^ expression to onset of ECM

Previously, we had noted a decreased number of activated CD8 T cells in inguinal LNs of PbA infected Cdh5-Cre D^b^ cKO mice (Fig. 3p). In order to further rule out peripheral immune perturbations and fully demonstrate that neuropathology was dependent on CD8 T cell interactions with MBMECs, we performed parabiosis (Fig. 7a). Surgery was performed to conjoin D^b^ expressing Cre- littermate controls with Cdh5-Cre D^b^ cKO mice in order to exchange peripheral blood and circulating immune cells. Three weeks of healing and acclimation were observed before both mice were I.P. injected with PbA to induce ECM (Fig. 7b). Our analysis revealed that all peripheral immune activation was similar in the spleen and inguinal LNs of parabionts after infection (Fig.7c-d). In the brain, no differences in CD45+ immune cell frequencies were observed and CD8 and CD4 T cell numbers mirrored that of the previous mouse cohorts with no difference between groups (Fig. 7e). Also consistent with prior cohorts, the Cdh5-Cre D^b^ cKO parabionts had decreased IFN-y expression by brain infiltrating CD8 T cells (Fig. 7e). We performed vascular permeability assays in our D^b^ expressing Cre- to Cdh5-Cre D^b^ cKO parabionts and determined that the vascular leakage phenotype was consistent with our previous data on D^b^ Cre- mice and had enhanced FITC-albumin leakage. In contrast, despite having circulating immune cells from D^b^ expressing Cre- mice, Cdh5-Cre D^b^ cKO mice resembled uninfected mice in that no vascular permeability was detectable, directly supporting our previous data (Fig. 7f). This observation was further supported by our results employing dual gadolinium enhanced T1-weighted MRI on the parabionts. We observed diffuse enhancement throughout the brain of the Cre- parabiont, indicating leakage. However, the conjoined Cdh5-Cre D^b^ cKO mice presented with no vascular leakage as demonstrated by lack of gadolinium-enhancement with T1-weighted MRI and retainment of FITC-albumin within vasculature (Fig. 7g). This data confirms that peripheral immunity was not impeded and that changes in infiltration and activation status are truly barrier and brain-specific and are impacted by class I restricted antigen presentation by MBMECs.

**Fig 7.**
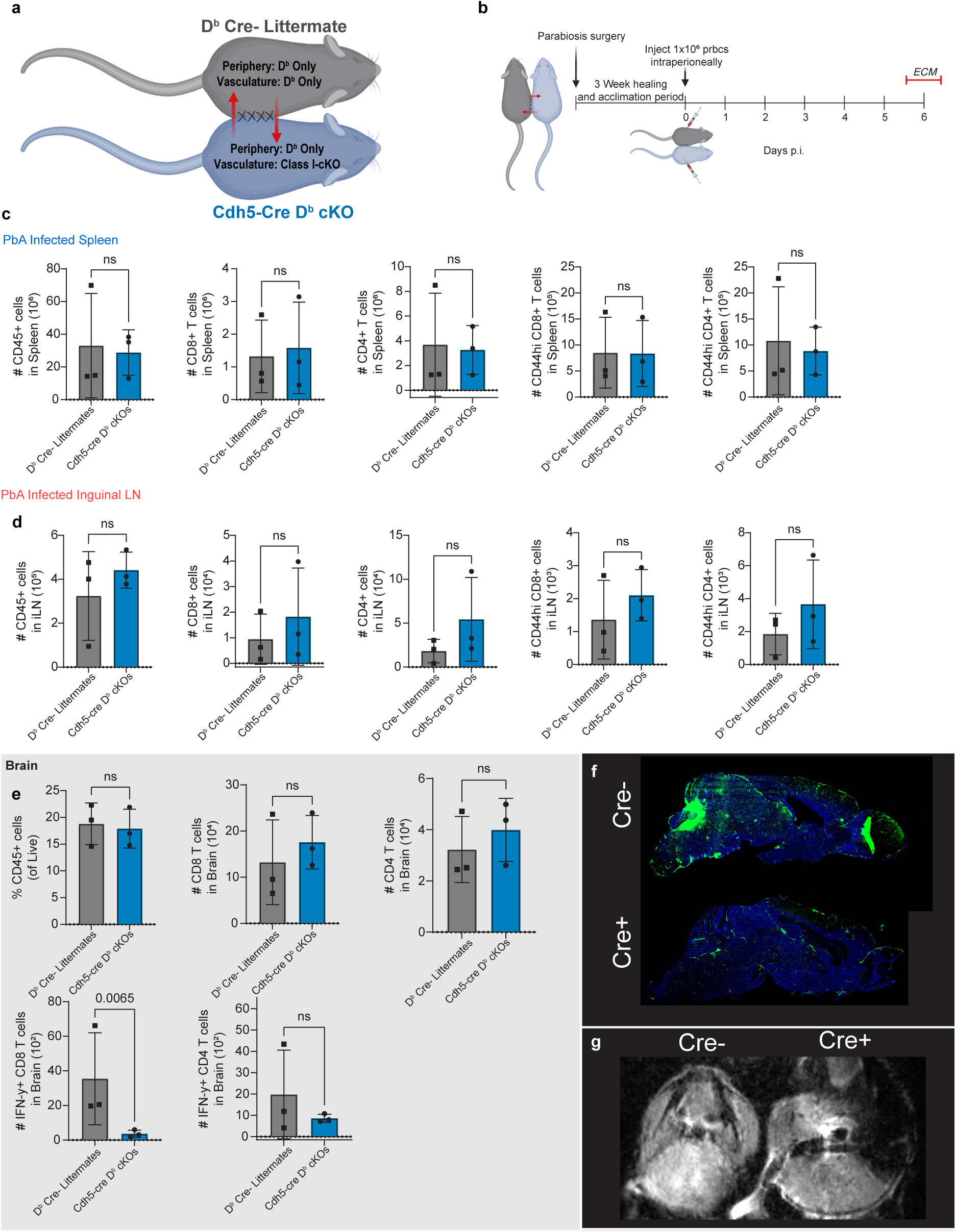
PbA infected parabionts sharing peripheral circulation demonstrate requirement of endothelial D^b^ expression for T cell responses and development of ECM. a) Schematic depicting parabiosis procedure and respective D^b^ Cre- littermate and Cdh5-cre D^b^ cKO mice. b) Timeline of experiment depicting parabiosis surgery, healing and acclimation time and PbA infection of parabionts. c) Absolute counts of total CD45+ immune cells, CD8 T cells, CD4 T cells and activated CD8s and CD4s in the spleens of parabionts day 6 of PbA infection. d) Absolute counts of total CD45+ immune cells, CD8 T cells, CD4 T cells and activated CD8s and CD4s in the iLNs of parabionts day 6 of PbA infection. e) Absolute counts of total CD45+ immune cells, CD8 T cells, CD4 T cells and IFN-y producing CD8s and CD4s in the brains of parabionts day 6 of PbA infection. f) FITC-albumin was IV injected one hour before euthanasia of PbA infected parabionts. D^b^ Cre- littermate and Cdh5-cre D^b^ cKO parabiont brains are visualized with FITC leakage in green. g) T1-weighted gadolinium enhanced dual MRI of parabiont pair was performed. Representative image is one axial slice with visualization of gadolinium enhancement. Data were analyzed with unpaired t test. Bar plots present the mean ± SD. All p-values are shown or ns for non-significant p-value *(p > 0.05)*.

### Differential Endothelial Class I expression alters the neuroinflammatory profile of the brain and targeted cell death during ECM

To define downstream genes discretely regulated by either K^b^ class I molecule or D^b^ class I molecule, we examined RNA transcriptomics data for differentially regulated genes in PbA infected wildtype C57BL/6, K^b^ Cre-, Cdh5-Cre K^b^ cKO, D^b^ Cre-, Cdh5-Cre D^b^ cKO, and uninfected controls of each group. We observed overlaps in gene expression that were shared between wildtype C57BL/6 mice (with K^b^ and D^b^) and K^b^ Cre- mice (K^b^ only) (Fig. 8a). Gene programs regulated by K^b^ were genes primarily expressed in anti-inflammatory A2 astrocytes, primed microglia, and cytokine and interferon signaling (Fig 8a and Supplementary Table 3). Cdh5-Cre K^b^ cKO mice had very few differentially regulated genes, similar to uninfected mice (Supplementary Figure 8a-c). We also noted distinctively differential expression of genes that overlap between wildtype (with K^b^ and D^b^) and D^b^ Cre- mice (D^b^ only) (Fig. 8a). Gene programs regulated by D^b^ Cre- mice were indicative of a pro-inflammatory A1 astrocytic response, stage 2 damage-associated microglia, and microglial neurodegenerative phenotype signatures^30, 31, 32, 33^. We also noted upregulation of genes associated with enhanced Ag-presentation and cytotoxic T cell responses in the brains of these animals, indicating a strong T cell response in these animals (Fig. 8a and Supplementary Table 3). Cdh5-Cre D^b^ cKO mice had very few differentially regulated genes (Supplementary Figure 8d-f). Overall, the differential gene expression observed between mice expressing distinct class I molecules further supports the independent roles each class I contributes to this pathology.

**Fig 8.**
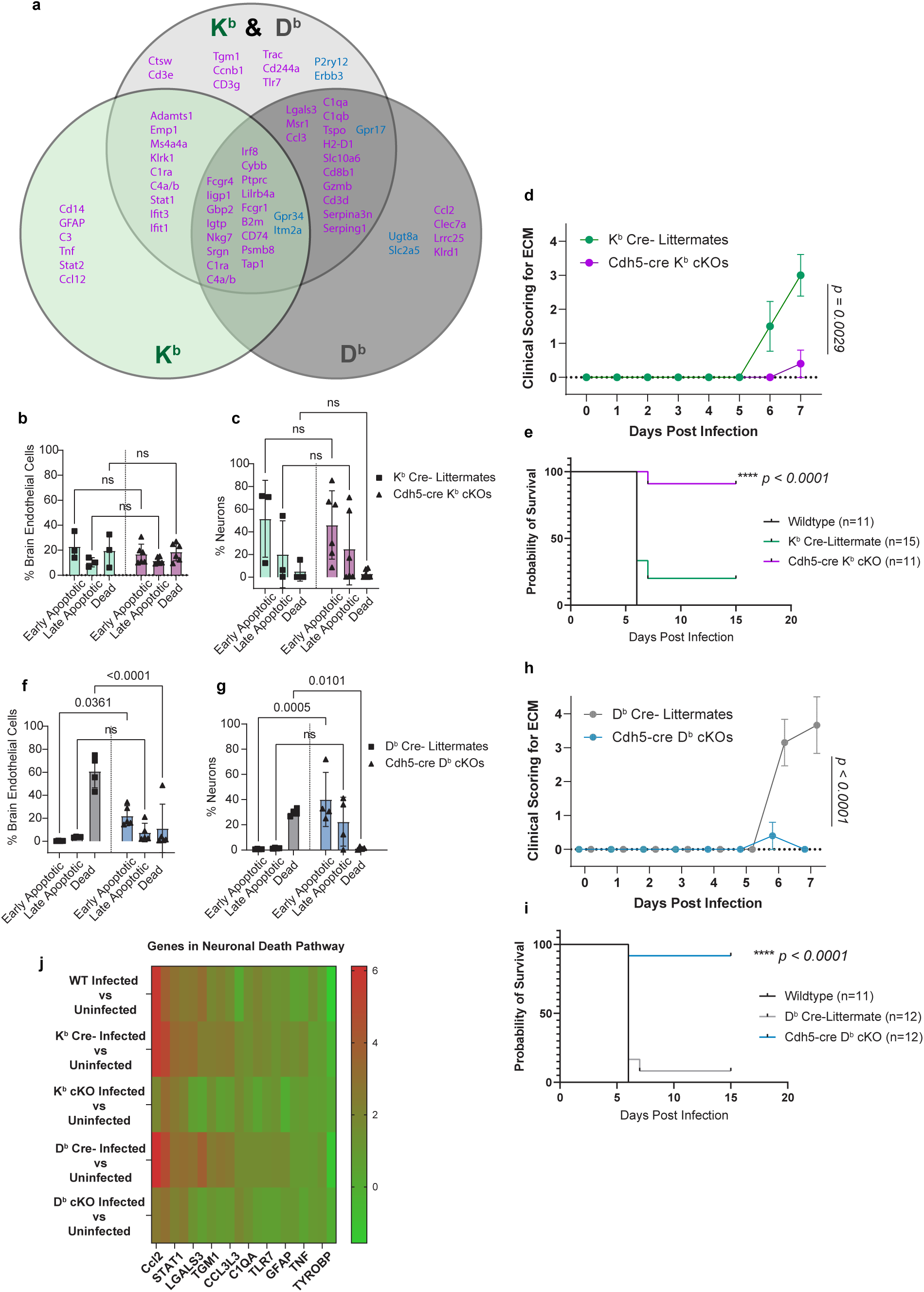
Discrete class I molecules differentially regulate cell death and neuroinflammation in the brain during ECM. Nanostring analysis was performed on whole brain isolated RNA of PbA infected C57BL/6, K^b^ and D^b^ Cre- littermates, Cdh5-cre K^b^ cKO, Cdh5-cre D^b^ cKO and uninfected controls of each group (n = 40, n = 4 mice/group). a) Venn diagrams of differentially expressed genes are visualized for comparison of differential class I regulation by wildtype mice, mice with only K^b^ and mice with only D^b^ expression. High-dimensional spectral flow cytometry was performed using a staining panel specific for stages of cell death. Frequencies of cells positive for markers of early apoptosis, late apoptosis and necrosis (late cell death) determined by annexin V staining vs viability stain gating. b) Frequencies of respective stages of cell death in Cre- littermates and Cdh5-cre K^b^ cKO expressed by brain endothelial cells on day 6 PbA infection. c) Frequencies of respective stages of cell death in Cre- littermates and Cdh5-cre K^b^ cKO expressed by mature neurons on day 6 PbA infection. d) Blinded clinical scoring was conducted daily on Cre- littermates and Cdh5-cre K^b^ cKO mice infected with PbA. e) Survival studies were performed on infected Cre- littermates and Cdh5-cre K^b^ cKO mice. Mice with K^b^ deletion had largely improved survival. f) Frequencies of respective stages of cell death in Cre- littermates and Cdh5-cre D^b^ cKO expressed by brain endothelial cells on day 6 PbA infection. g) Frequencies of respective stages of cell death in Cre- littermates and Cdh5-cre D^b^ cKO expressed by mature neurons on day 6 PbA infection. h) Blinded clinical scoring was conducted daily on Cre- littermates and Cdh5-cre D^b^ cKO mice infected with PbA. i) Survival studies were performed on infected Cre- littermates and Cdh5-cre D^b^ cKO mice. Mice with D^b^ deletion had vastly improved survival. j) IPA analysis was performed, and gene Expression Log Ration data was plotted in heat map to visualize changes in expression between groups. Data were analyzed with unpaired t test or 2way ANOVA with multiple comparisons. For survival, Kaplan Meier curves were plotted, and Log-rank (Mantel-Cox) curve comparison was performed. Scatter plots and bar plots present the mean ± SD. All p-values are shown or ns for non-significant p-value *(p > 0.05)*.

We next wanted to determine what cells are being targeted by infiltrating CD8 T cells during PbA infection. We infected mice with PbA and harvested tissue early on day 6 before all mice were moribund. Brains were harvested to stain with a panel of antibodies to detect stages of cell death. In K^b^ expressing Cre- littermates, we observed no significant increases in cell death of MBMECs on day 6 of infection and no differences when K^b^ is deleted on vasculature (Fig. 8b). When assessing neuronal cell death however, there were increased frequencies of cells with early apoptotic markers, but complete neuronal cell death was not detectable by flow at this time. There were no differences between K^b^ expressing Cre- littermates and Cdh5-Cre K^b^ cKO mice (Fig. 8c). The low levels of endothelial and neuronal cell death may be attributed to the fact that pathology is delayed in K^b^ mice in comparison to D^b^ mice. The K^b^ Cre- mice have average clinical scores of ∼1.5 on day 6 for ECM neurological signs, whereas D^b^ Cre- mice have average clinical scores of >3 day on 6 (Fig. 8d and 8h). This delay in pathology may also translate to a delay of neuronal and/or endothelial cell targeting as well. To examine this, we performed another experiment in which all mice were allowed to progress to humane endpoint. Transcriptomics analysis revealed that neurons do activate cell death pathways at this late timepoint in K^b^ Cre- littermates (Fig. 8j). No endothelial cell death pathway activation was noted. Cdh5-Cre K^b^ cKO mice had significantly improved clinical scores when K^b^ on vasculature was deleted (Fig. 8d). In extended survival studies, we observed nearly complete survival of Cdh5-Cre K^b^ cKO mice in comparison to Cre- littermate controls (Fig. 8e).

When assessing D^b^ mouse cohorts at day 6, we observed a striking increase in MBMEC cell death in Cre- littermates that have D^b^ in comparison to Cdh5-Cre D^b^ cKO mice (Fig. 8f). A significant increase in neuronal cell death was observed in D^b^ expressing Cre- littermates in comparison to Cdh5-Cre D^b^ cKO mice (Fig. 8g). Similar to our observations in K^b^, we observed increased frequency of neurons with early apoptotic markers in the Cdh5-Cre D^b^ cKO mice, but not late-stage neuronal death (Fig. 8g). These data were in agreement with transcriptomics data showing activation of cell death pathways only in D^b^ Cre- mice (Fig. 8j). In an extended survival study, mice with deletion of D^b^ on vasculature presented with striking improvement in clinical scores and overall survival (Fig. 8h-i). Overall, neuronal cell death related gene expression was increased in infected wildtype C57BL/6, K^b^ expressing Cre- mice and D^b^ expressing Cre- mice (Fig. 8j). These studies demonstrate that class I molecule expression by endothelium is required for the onset of neuronal cell death in the ECM model and that MBMECs undergo cell death late in pathology following D^b^ MHC Class I-specific-CD8 interactions at vasculature.

## Discussion

The role of Ag-presentation by brain endothelium and the effect on the CD8 T cell effector response as a putative mechanism of CM have been highly debated in the cerebral malaria field^3, 25, 34^. However, the role of individual class I molecule presentation by MBMECs to brain infiltrating CD8 T cells has never been directly tested *in vivo*. In this study, we demonstrate that targeted deletion of either K^b^ or D^b^ class I molecule on endothelium results in reduced T cell interactions at brain vasculature and subsequently attenuated BBB disruption and neuropathology. Comparisons to human data supports these findings, as well as other recent studies which highlighted the importance of CD8 T cell interactions at the vasculature in the development of CM and ECM^11, 17^. Overall, peripheral infection and all immune responses develop normally in our mice, with the exception of mild decrease in CD8 T cell activation marker CD44 at baseline in non-brain draining inguinal lymph nodes in Cdh5-Cre K^b^ cKO mice. However, this difference resolves during infection. In Cdh5-Cre D^b^ cKO animals there was also a slight decrease in the CD8 T cell activation marker CD44 in non-brain draining inguinal lymph nodes during PbA infection. We addressed the potential effects these differences could pose in ECM by performing parabiosis, which efficiently equalizes peripheral immune responses. Endothelial cell-specific deletion of class I molecule was still protective, effectively ruling out potential peripheral immune perturbations as having a role in protection from ECM.

CD8 T cells have been observed to interact with brain vasculature during neuroinflammation^17, 26, 35, 36, 37^. However, few of these molecular interactions have been successfully interrogated. The most well-characterized of these interactions is the CD8 T cell slowing, rolling, and arrest on brain endothelium^38, 39, 40, 41, 42^. Using intravital microscopy on live mice 6-7 dpi with PbA, we observe high numbers of T cells in both K^b^ and D^b^ expressing Cre- littermates exhibiting slowing, rolling and arresting. In contrast, we observed a drop in these T cell interactions with brain vasculature with the loss of either K^b^ or D^b^ class I expression by endothelial cells. The novelty of these findings, however, lies in the fact that the adhesion molecule expression is completely unchanged between Cre- and Cre+ animals in the K^b^ model. This implies that the complete loss of T cell interactions we visualize by 2-photon are dependent on the T cell-K^b^ class I interaction and not adhesion molecules. In D^b^ expressing Cre- mice, we observe upregulation of P-selectin on brain endothelium. Surprisingly, despite their essential role in T-cell rolling in vasculature, absence of E or P-selectin in mice have been experimentally shown to be unnecessary for T-cell entry into the CNS, suggesting that T cell rolling is not required for T-cell migration across inflamed BBB ^39, 40, 41, 42^. However, most studies focusing on diapedesis interrogated CD4 T cell interactions. ICAM-1 and VCAM-1 on endothelium was not altered when class I molecules were ablated on vascular endothelial cells. Interestingly, when either class I molecule was deleted on endothelium, expression of MHC Class II molecules was found to be elevated. This may be a compensatory mechanism employed by endothelial cells in order to establish a more robust CD4 T cell response when the CD8 T cell response is impaired^43^. The ability of brain endothelium to shift the T cell effector response in the brain is in agreement with several studies which propose that endothelial cell Ag-presentation may have important immunomodulatory functions^35, 36, 37, 44^.

Observing that deletion of both K^b^ and D^b^ class I molecule on endothelial cells resulted in loss of T cell interactions with brain vasculature, we expected this may result in loss of CD8 T cell infiltration. Interestingly, we observed differential CD8 T cell infiltration into brain parenchyma. In Cdh5-Cre K^b^ cKO mice, we saw a reduction of CD8 T cells infiltrating the brain. However, in Cdh5-Cre D^b^ cKO mice, we saw no changes in CD8 T cell infiltration, indicating that overall CD8 T cell infiltration and D^b^-restriction may be uncoupled in this model. Although the deletion of D^b^ does not seem to affect overall infiltration of CD8 T cells, lack of D^b^ does alter effector status. It is known that PbA strain of *Plasmodium* causes brain specific sequestration of prbcs and activation of brain microvasculature^25^. However, work describing the PbA antigenic D^b^-restricted peptide SQLLNAKYL showed that when mice are infected with strains of *Plasmodium* that do not cause adherence of prbcs to brain endothelium, CD8 T cells still entered the brain but were not activated, resulting in absence of neurological symptoms in these animals^45^. This supports our findings that when *Plasmodium* initiates endothelial cell Ag- presentation, D^b^ is required for activation of CD8 T cells, but not infiltration during infection. Furthermore, CD8 T cell TCR binding to cognate peptide presented by both H-2K^b^ and H-2D^b^ on brain endothelium resulted in IFN-γ production by infiltrating CD8 T cells, which in turn, caused differentially regulated cell death and mortality of the animal. This is supported by other works showing that IFN-y production induces a cascade to recruit more CD8 T cells to the brain^43, 45, 46^. Overall, we demonstrated that endothelial K^b^ class I molecule interactions with CD8 T cells is important for infiltration and activation, while endothelial D^b^ class I molecule interactions activate CD8 T cells but does not influence diapedesis into parenchyma.

Previous work from our lab and others has hinted at the importance of antigen-specificity of these infiltrating CD8 T cells in ECM neuropathology^16, 25, 45, 47^. However, definitive support for this has been elusive and incomplete^17, 45^ ^48^. Studies with transgenic parasites bearing model epitopes are unable to recapitulate the complexity of the diverse CD8 T cell responses against the wide array of native malaria Ags seen in mice and humans^49^. The epitopes that are known have been shown to elicit variable CD8 T cell responses^49^. Although epitope discovery continues to be an active area of research, there are currently only 83 known T cell epitopes for PbA out of what is projected to be many thousands^35, 45, 50^. Importantly, a role for BBB damage-associated molecular patterns (DAMPs) or parenchymal brain draining DAMPs in eliciting CD8 T cell responses, have also not been ruled out^25, 34, 49, 50^. To date, our novel mouse models are the first models capable of distinguishing between K^b^ and D^b^ class I restricted Ags specifically presented by brain endothelial cells for interrogation of ECM pathogenesis. To assess the role of cognate TCR-MHC interactions in CD8 T cell infiltration in this pathology, we used the well-established surrogate marker strategy to track Ag-experienced CD8 T cells^28^. We saw no differences in Ag-experienced CD8 T cells in the meninges or cervical LNs in either model. However, Ag-experienced CD11a+ CD8 T cells were reduced specifically in brain parenchyma when lacking K^b^ or D^b^ on vasculature^28, 51^. We also employed IV labeling of these infiltrating Ag-experienced CD8 T cells 3 minutes prior to harvesting tissue. The caveat of this work is that IV labeling may increase in parenchyma with increased vascular permeability, although this does not seem to be the case in the K^b^ model. We saw that infiltration is much higher in K^b^ Cre- littermates than in Cdh5-Cre K^b^ cKO mice at day 6, and that most CD8 T cells had previously infiltrated over time prior to IV administration of labeling antibody. In the D^b^ model, however, we see comparable amounts of IV labeled and IV negative cells in the brain indicating that a large influx of CD8 T cells begins day 6 of pathology. These data suggest that K^b^ Ag-presentation by brain endothelium tightly regulates activation and entry. However, D^b^ Ag-presentation by brain endothelium does not employ as strict of regulation of CD8 T cell infiltration at the barrier. This may mean that another predominant interaction, by a currently undefined cell type or chemokine gradient, may be initiating this brain infiltration. This other interaction, however, does not sufficiently activate the T cell as is seen when brain endothelium presents on D^b^. Overall, these data suggest that CD8 T cell activation and infiltration are influenced differentially in the D^b^ model during ECM.

The chemokine receptor expression of activated T cells shows dynamic plasticity. Several factors, including strength of antigenic signals, tissue-specific imprinting by priming DCs, or cytokine milieu can influence the trafficking of T cells^52^. Both CXCR3 and CCR7 are receptors on CD8 T cells shown to be necessary for recruitment into the brain to establish ECM^24, 25, 26^. During ECM, olfactory bulb and brainstem are physically and functionally damaged, with high prbc and leukocyte sequestration in these regions, which coincides with vascular pathology and neuronal cell death^17, 27^. Specifically, the olfactory region has been shown to highly upregulate CCL21, while brain endothelial cells highly express CXCL9 and neurons highly express CXCL10^24, 27, 53^. Blocking their receptors CCR7 and CXCR3 results in decreased CD8 T cell activation and recruitment to the CNS, respectively, as well as prolonging survival during ECM^27, 54^. To support that T cells were sufficiently primed in the periphery and imprinted for CNS-specific trafficking we examined chemokine receptor expression on CD8 T cells trafficking to the brain. In the brain, frequency of CD8 T cells expressing CXCR3 and CCR7, and MFIs of receptor expression were unchanged between Cre- and cKO animals in both models; thereby reinforcing brain-specificity. The majority of data on CD8 T cell infiltration in ECM focuses on the cortex, as this is the brain region most heavily infiltrated by CD8 T cells during this pathology^55^. However, no prior reports have shown regional infiltration of areas affected by vascular permeability. In this work we not only show regional CD8 infiltration to areas with highest brain pathology, but that CD8 T cells are also present in regions without vascular leakage, a topic previously debated^17^.

CNS vascular leakage was prevalent in both K^b^ and D^b^ class I molecule expressing mice. However, the presentation was distinct. In comparison to wildtype C57BL/6 animals, in which both class I molecules are expressed, we saw less leakage in the midbrain of K^b^ expressing animals, although there was still significant damage and leakage to olfactory and brainstem regions. Alternatively, D^b^ expressing animals presented with worsened pathology, more closely resembling that of the wildtype C57BL/6 mice which express both class I molecules. Importantly, when either K^b^ or D^b^ were ablated specifically from vasculature, there was complete rescue of vascular permeability. This finding is highly novel and strongly suggests that antigens are being presented by either K^b^ or D^b^ class I molecules, specifically by brain endothelial cells, to elicit distinct neuropathology associated with ECM. We also show that Claudin-5 is altered in regions of permeability in Cre- animals, a known hallmark of vascular pathology in this disease^16, 17^. However, animals lacking class I on vasculature preserved normal Claudin-5 expression in the regions of interest.

We observed that K^b^ and D^b^ Ag-specific activation of recently infiltrated CD8 T cells expressing IFN-y correlates with differential vascular permeability in ECM. We therefore wanted to interrogate whether K^b^ and D^b^ class I molecules produced differential activation of neuroinflammatory pathways. We did observe distinct patterns of neuroinflammation, with differential activation of astrocytes and changes in microglial and T cell activation states. This may explain why D^b^ restricted Ag-presentation on endothelium, though resulting in later infiltration of Ag-experienced CD8 T cells than K^b^ restriction on day 6, still resulted in stronger T cell signaling, more cell death and worsened vascular leakage and survival at endpoint. Taken together, these data indicate that discrete class I molecules on brain endothelium have immunomodulatory capacity, capable of altering the neuroinflammatory profile of the brain during ECM.

Knowing that K^b^-restriction and D^b^-restriction incite differential neuroinflammatory profiles in the brain during PbA infection, we next sought to determine whether cellular targets of the CD8 T cells were also distinct. We demonstrated that 6 dpi, mice with K^b^ class I molecules displayed no markers of apoptosis or cell death in vascular endothelium. However, neurons had early apoptotic markers and transcriptomic data supported that neuronal death is induced at endpoint. When K^b^ was deleted from vasculature, endothelial cells were unchanged. However, neurons no longer acquired the gene signature indicative of neuronal death. This data suggests that CD8 T cell interactions with K^b^-restricted Ag-presenting endothelial cells are not required for targeted killing of MBMECs but are supportive of apoptotic neuronal cell death late in pathology. The K^b^-CD8 T cell interaction has been shown to modulate infiltration and activation of CD8 T cells and may be involved in CNS reactivation or serve an imprinting role which may instruct CD8 T cells to exact their effector functions in other ways or on other cell types. When assessing D^b^ Cre- littermates, we observed a large percentage of cell death of MBMECs which was ameliorated when D^b^ was deleted on vasculature. There was also an increased percentage of cell death seen in neurons in D^b^ expressing Cre- animals which was either resolved or delayed when D^b^ was lost on vasculature. This data demonstrates a requirement for D^b^-restricted Ag-presentation to CD8 T cells by the vasculature in order to enact an effector response resulting in killing of both MBMECs and ultimately neuronal cell death. The extent to which neuronal cell death is the result of direct killing by CD8 T cells, or through alternative mechanisms has yet to be explored in these models. In previous work which detailed CD8 T cell responses during ECM, they demonstrated that K^b^ and D^b^ are capable of presenting similar peptides, though with very different binding affinities^49^. It is also known that receptor affinities control effector responses and T cell polyfunctionality^45, 56, 57, 58, 59^. This is consistent with our data showing that endothelial cells die when D^b^ presents, but not when K^b^ presents, dependent upon the differential peptides they display as well as the strength of their interactions. These results prove to be highly innovative as this is the first time that *in vivo* evidence has directly shown the effects of K^b^ or D^b^ restricted Ag-presentation by brain endothelial cells during ECM. Follow-up investigations should address direct mechanisms of cell death in more detail. Alterations of cytokine milieu, the impact of vascular leakage and blood-derived factors on neuroinflammation and CD8 T cell responsiveness still must be addressed.

In conclusion, our studies provide the first *in vivo* interrogation of the role of Ag-presentation by individual class I molecules by MBMECs in developing ECM pathology. We have dissected the contribution of both K^b^ and D^b^ to the onset of neuropathology and provide evidence to support that K^b^-restricted Ag engagement on endothelium plays a critical role in CD8 T cell vascular adhesion, arrest, infiltration, and activation. These events in turn contribute to blood-brain barrier tight junction downregulation, vascular leakage, and neuronal death. Meanwhile, D^b^-restricted Ag engagement on endothelium plays a role in vascular adhesion and arrest but is inconsequential for CD8 T cell infiltration into brain parenchyma. However, the interaction is important for CD8 T cell activation and IFN-y secretion which is associated with heightened CD8 T cell motility and toxicity, tight junction downregulation, and vascular permeability^25, 29, 46, 60^. Strikingly, we show that only CD8 TCR recognition of D^b^-restricted antigens by brain vasculature results in killing of MBMECs and subsequent neuronal cell death. We contend that these insights into the pathogenesis of cerebral malaria will provide rationale for therapeutic targeting of these interactions. Anti-malarial drugs can serve to lessen parasite burden, but show no improvement in neurological symptoms once there is brain involvement^17^. We therefore propose that patients with CM may benefit from endothelial-specific therapeutics which serve to specifically blockade T cell-class I interaction in addition to standard-of-care anti-malarial drugs.

## Methods

### Animals

Male and female C57BL/6J WT (stock no. 000664) and Cdh5-Cre mice (stock no. 033055) were purchased from Jackson Laboratory. H-2K^b^ LoxP and H-2D^b^ LoxP mice were generated by our program as previously described, in the C57BL/6 background ^61, 62^. H-2K^b^ LoxP and H-2D^b^ LoxP mice were crossed with Cdh5-Cre mice to generate Cdh5-Cre K^b^ cKO and Cdh5-Cre D^b^ cKO mice that express Cre under the *Cdh5* promoter and undergo endothelial-specific targeted gene deletion of H-2K^b^ and H-2D^b^ molecules. Mice were then bred and maintained under specific pathogen free conditions at the Mayo Clinic Animal Facility. In the studies, 15–20-week-old mice were used for all experiments with appropriate age and sex-matched controls. All experiments were conducted in accordance with guidelines from the National Institutes of Health and under the approval of the Mayo Clinic Institutional Animal Care and Use Committee.

### Plasmodium infection

*Plasmodium berghei* ANKA (PbA) stocks were generously provided by J. Harty at the University of Iowa and S. Hamilton Hart at the University of Minnesota Medical School. PbA infection was induced in 15–20-week-old mice via intraperitoneal injection of ∼10^6^ parasitized red blood cells on day 0. Mice were monitored twice daily until onset of clinical signs were observed (days 6-8 post infection), and then continuously. Blinded scores were obtained daily, scoring signs of neurological impairment until endpoint. Mice were ranked for clinical signs of ECM adapted from Hortle et al^63^: 0, no clinical symptoms; 1, reduced movement and limp/flaccid tail; 2, rapid breathing and transient hunching posture; 3, hunched, ruffled fur and loss strength in four limb hang test; 4, severe disease (ataxia, seizures, coma); and 5, death. Moribund animals were scored 4.5 and humanely euthanized. Mice were classified as having ECM if displaying symptoms between days 6-8 post-infection. Peripheral parasitemia was assessed using thin blood smears and Geimsa staining as outlined on the Centers for Disease Control website and as previously described^64^.

### Acute TMEV and TMEV-OVA Infection

The Daniel’s strain of TMEV was prepared as previously described^62^. TMEV-XhoI-OVA8 (TMEV-OVA) was generated and prepared by our group as previously described^23^. At 10-12 weeks of age, mice were anesthetized with 1-2% isoflurane and infected intracranially (i.c.) with 2 x 10^6^ PFU (10µL) of the Daniel’s strain of TMEV, or 2 x 10^5^ PFU of TMEV-OVA. Virus was delivered to the right hemisphere of the brain via automatic 1 mL Hamilton syringe (Hamilton Company, Reno, NV). Mice were euthanized for high-dimensional flow cytometry of tissues at 7-days post infection (dpi).

#### Magnetic resonance imaging (MRI)

MRI capture and analysis was conducted as previously reported using Bruker Avance II 7 Tesla or Bruker Avance 16.4 Tesla vertical bore small animal MRI system (Bruker Biospin, Billerica, MA)^65^. Gadolinium (Multihance-Bracco) was administered intraperitoneally (100µL of 100mg/kg) ∼15 minutes before scans. 25 mm and 34 mm volumetric coils were utilized. Animals were anesthetized with continuous isoflurane administered through nose cone. Respiration and heart rate were monitored with an MRI compatible vitals monitoring system (model 1030; SA Instruments, Stony Brook, NY). After axial and sagittal tripilot positioning scans, isotropic 3D T1-weighted scans were employed to detect gadolinium, with an approximate scan time of 30 minutes. Maximum intensity projection scans were performed to detect gadolinium, with an approximate scan time of 10 minutes. Scoring was performed by two independent observers, blinded to genotype with <10% intra-observer variation found. Analyze12.0 software (Biomedical Imaging Resource; Mayo Clinic, Rochester, MN) was employed to analyze T1 data and to generate 3D maps of gadolinium enhancement. Maximum intensity projection data was analyzed using Fiji Is Just ImageJ (FIJI) 2.3.0; Java 1.8.0_172.

#### Immunofluorescence

Brains used for immunofluorescence staining were either fresh frozen or paraformaldehyde fixed following mouse euthanasia and transcardial perfusion with ice-cold PBS. Brains harvested after 2-photon imaging and FITC-albumin injection were harvested without perfusion and paraformaldehyde fixed. All brain tissues were embedded in Tissue-Tek OCT Compound, frozen on dry ice and sliced on cryostat with the exception of FITC-albumin injected brains which were embedded in 4% agarose and sliced with vibratome. All brain slices were mounted on SuperFrost Plus microscope slides and stored at 4 degrees.

#### Single-cell suspension

Mice were euthanized via isoflurane per IACUC guidelines and transcardially perfused with ice-cold PBS. Brains and meninges were harvested and processed as previously described^66, 67^, via dounce homogenization and 30% Percoll gradient or enzymatically digested using mouse tumor dissociation kit and gentleMACS™ Octo Dissociator (Miltenyi Biotec). Spleens and lymph nodes were either digested with Collagenase IV/DNAse or not digested before mechanical disruption. Lymph nodes were dissociated using frosted glass slides and spleens were dissociated through 70 µM filter. Thymus samples were processed using mechanical disruption through 70 µM filter. Blood was collected (100µl) into 1 mL heparin (100U/mL). Spleens underwent ACK RBC lysis for 3 minutes and blood samples for 1minute. All samples were filtered using 70 µM filters before counting, staining and running on flow cytometer.

#### Flow cytometry

After resuspension in PBS, cells were either first stained with either Zombie UV or Zombie NIR fixable viability dyes (BioLegend) (1:1000) for 20 minutes at room temperature in the dark or Ghost Red 780 Viability dye (Tonbo) was added to surface stain master mix. Samples underwent Fc-receptor block and staining for 30 minutes at 4 degrees with surface staining antibodies. Panels requiring intracellular staining were then fixed and permeabilized using Cyto-Fast™ Fix/Perm Buffer Set (BioLegend), and then stained with intracellular antibodies for 30 minutes at room temperature in the dark before washing. All flow panel staining was performed with this method with the exception of Annexin staining, which required staining in Annexin V Binding Buffer (Invitrogen) for 15 minutes at room temperature prior to fixation and permeabilization and intracellular staining. The antibodies that are specific for surface markers included CD103 Brilliant Violet 421 (121422; BioLegend), NK1.1 Pac Blue (108722; BioLegend), B220 Pac Blue (103227; BioLegend), CD62L Brilliant Violet 480 (746726; BD Biosciences), CD11b VioGreen (130-113-239; Miltenyi Biotec), CD44 Brilliant Violet 570 (103037; BioLegend), CD80 Brilliant Violet 605 (104729; BioLegend), Ly6C Brilliant Violet 650 (128049; BioLegend), H-2K^b^/H-2D^b^ Brilliant Violet 750 (746869; BD Biosciences), CCR7 Brilliant Violet 785 (120127; BioLegend), I-A/I-E BB515 (565254; BD Biosciences), CD8a Spark Blue 550 (100780; BioLegend), CD45 PerCP (103130; BioLegend), Ly6G PerCP-eFluor 710 (46-9668-82; Invitrogen), OX40L PE-Dazzle 594 (108816; BioLegend), CD40 PE-Cy5 (124618; BioLegend), CD31 PE-Cy7 (25-0311-82; Invitrogen), CD3 Spark NIR 685 (100262; BioLegend), CD11c Alexa Fluor 700 (117320; BioLegend), CD4 APC-Fire 750 (100568; BioLegend), CD25 Brilliant Violet 421 (102034; BioLegend), TCRB APC (20-5961-U100; Tonbo biosciences), c-kit Brilliant Violet 786 (564012; BD Biosciences), CD69 Brilliant Violet 650 (740460; BD Biosciences), Ly51 FITC (108305; BioLegend), EpCAM PE-Cy7 (118216; BioLegend), CD8a Pac Blue (563898; BD Biosciences), CD4 APC (100516; BioLegend), CD11a PE (141006; BioLegend), CD16/CD32 (553141; BD Pharmingen), Thy1.2 Alexa Fluor 488 (105316; BioLegend), I-A/I-E Pac Blue (107620; BioLegend), CD8a Brilliant Violet 510 (100751; BioLegend), CD44 Brilliant Violet 650 (740455; BD Biosciences), Ly6G Brilliant Violet 711 (46-9668-82; Invitrogen), CD4 Brilliant Violet 786 (563727; BD Biosciences), Ly6C PerCP (128027; BioLegend), CD45 PE-CE594 (562420; BD Horizon), CD11b PE-Cy5 (55-0112-UlOO; Tonbo biosciences), TCRB PE-Cy7 (60-5961-U100; Tonbo biosciences), B220 Spark NIR 685 (103268; BioLegend), CD62L APC-Fire 750 (104450; BioLegend), CD11c Brilliant Violet 605 (117333; BioLegend), Annexin V(25-8103-72; Invitrogen), B220 APC-Fire 810 (103277; BioLegend), CD26 Brilliant Violet 605 (745125; BD Biosciences), CD80 Brilliant Violet 711 (104743; BioLegend), I-A/I-E Brilliant Violet 786 (743875; BD Biosciences), CD8a Brilliant Violet 421 (563898; BD Biosciences), CD31 APC-Fire 750 (102528; BioLegend), CD11a PE-Cy7 (101122; BioLegend), CD31 PE-Dazzle (102526; BioLegend), CXCR3 Brilliant Violet 421 (126521; BioLegend), CD106 Alexa Fluor 647 (105712; BioLegend), CD62P PE (148306; BioLegend), CD54 Brilliant Violet 711 (116143; BioLegend), CD8 Brilliant Violet 570 (100740; BioLegend), Zombie NIR viability dye (Cat. #423105; BioLegend), Zombie UV (423107; BioLegend). To assess cytokine production cells were stained with intracellular markers after fixation and permeabilization: NeuN APC (100920; Novus), IFNy Brilliant Violet 785 (563773; BD Biosciences), IFNy APC (20-7311-U100; Tonbo biosciences) IL-12 PE (505204; BioLegend), IFNy PE (50-7311-U100; Tonbo Biosciences). In noted experiments, intravenous injection of 2ug of Thy1.2 Alexa Fluor 488 were administered 3 min. before collecting tissues to label T cells in vasculature. All samples were acquired on Cytek Aurora spectral flow cytometer (Cytek, Fremont, CA). All analysis was performed using FlowJo software version 10.8.1(FlowJo LLC, Ashland, OR).

#### *In vivo* two-photon microscopy

Mice were intravenously injected with 2 µg of Thy1.2 Alexa Fluor 488 just prior to cranial window surgery, approximately 30 min prior to imaging. Cranial windows were implanted as described ^68 69^. In brief, under isoflurane anesthesia (3% induction, 1.5–2% maintenance), a circular craniotomy (<5 mm diameter) was made over somatosensory cortex with the center at about -2.5 posterior and +2 lateral to bregma. A circular glass coverslip (5 mm diameter, Warner) was secured over the craniotomy using dental cement (Tetric EvoFlow). A four-point headbar (NeuroTar) was secured over the window using dental cement. Mice were injected intravenously with intravascular dye 70,000 MW Dextran-Texas Red (Invitrogen, D1830) 5 minutes before imaging. Alexa Fluor 488 (T cells) and Dextran-Texas Red (vessels) were excited by a tunable Ti:Sapphire Mai Tai DeepSee laser (Spectra Physics, CA, USA) at 940 nm and the emission was collected at 525/50 and 630/75 nm, respectively (U-MF2, Olympus, Japan). An Olympus LUMPLFLN 40x water-immersion objective was used to acquire image stacks. A 30-µm stack with 2 µm z-resolution and 512 × 512 pixels were acquired every minute for a period of 10-30 min. Images were analyzed by Fiji ImageJ ^70^. To analyze T cell velocity, each 30-µm stack was first aligned along the z axis using the StackReg plugin (Rigid Body) and Z-project into one image. Then images of every minute were stacked and aligned using StackReg plugin (translation). The T cell velocity was measured using the Manual Tracking plugin tracing for 10 min per field. The arrest coefficient is the percentage of each track that a cell is immobile (instantaneous velocity <2 µm/min) ^71^.

#### Parabiosis

Procedure was performed as previously described. In brief, two female age-matched mice of similar weight were selected. Mice were either littermates or co-housed for two weeks prior to surgery. Anesthesia was induced and maintained with Ketamine/Xylazine (100/10 mg/kg; maintenance dose 30/3 mg/kg). Ophthalmic ointment was placed on eyes. Mice are tested for reflex responses to ensure anesthetic depth. For analgesia during surgery, we administered Carprofen and Buprenorphine-SR subcutaneously (5 mg/kg and 1 mg/kg). Animals were shaved on the flank and prepped for surgery with Betadine swabs and sterile drapes. Using a sharp scissor, superficial epidermal incision was continuously made along the flank starting at 0.5 cm rostral of olecranon to 0.5 cm superior to patella. Skin absorbable 3-0 silk suture were used to join elbow and knee joints (Ethicon Inc, Somerville, NJ). Simple continuous suture was performed with absorbable 4-0 vicryl suture (Ethicon Inc, Somerville, NJ) for closer of incision. Post-operatively we administer 1 ml of 0.9% NaCl fluids subcutaneous, analgesics Carprofen intraperitoneally and Buprenorphine SR subcutaneously. Mice were placed on heating pad and observed for a minimum of 3 hours after surgery. Prophylactically, mice were treated with antibiotic Enrofloxacin (50 mg/kg) 10 days post-operatively. Parabiont pairs underwent a minimum of 3 weeks of healing and acclimation time before experimental use. Procedure was performed according to the IACUC guidelines at Mayo Clinic.

#### Nanostring

Brain dissections were subjected to RNA extraction (Qiagen RNAeasy Mini Kit Cat. No. 74104), followed by nanodrop concentration and purity analysis. RNA samples were applied to NanoString nCounter (NanoString Technologies, Seattle, WA) digital RNA quantification, using the predesigned mouse Glial Profiling Panel codeset. The Glial Profiling Panel reagents were provided by Nanostring. Reporter CodeSet Lot#RC8131X1. Capture Codeset Lot#CP8131X1. Nanostring analysis was performed by the Thompson Lab at the Mayo Clinic Comprehensive Cancer Center at Mayo Clinic Florida. Briefly, 100 ng of total RNA was hybridized to capture-reporter probe codesets and immobilized on NanoString cartridges. Excess RNA and probes were removed, and digital barcodes were counted. NanoString nCounter analysis software (v4.0) was used for counting, quality control, and normalization. Transcript counts were normalized according to NanoString’s recommendations. Normalization parameters were applied with normalization cutoff of 10. All samples were quality checked for quality and concentration. 13 housekeeping genes, 6 positive control genes, and 8 negative control genes were utilized for normalization.

### Human brain microvascular endothelial cells

Human brain endothelial cell microarray data available through public repository at Gene Expression Omnibus (GEO), accession number GSE9861. Outlined in the publication by Tripathi et al, HBMECs were isolated from human brain specimens and characterized as described previously^19, 72^. In brief, HBMECs (passage < 6) from single adult donor were used for the biologic replicate experiments. For microarray experiments, HBMECs were grown in collagen-coated 6-well plates until cells achieved about 80% to 90% confluence. The medium was then changed to RPMI 1640 with 5% fetal bovine serum for 24 hours before coincubation with Pf-IRBCs, at which time HBMECs are confluent. Endothelial cells mycoplasma infection negative (Vector Laboratories).

#### Gene Expression Analyses

Human microarray data with accession number GSE9861^19^ was utilized via Gene Expression Omnibus (GEO), a public functional genomics data repository supporting MIAME-compliant data submissions. Genes from human and mouse datasets were identified to be differentially expressed from each in comparison to a control group using either Ingenuity Pathway Analysis software (Qiagen), or Geo2R^73^. Gene set enrichment analysis (GSEA) was also performed on human data^74^.

#### Statistics

When comparing results from two groups, an unpaired Student’s t-test was used. When comparing results from two groups of paired mice, a paired Student’s t-test was used. When comparing results from three groups, a one-way analysis of variance (ANOVA) using Tukey’s post-hoc multiple comparisons test was used. A two-way ANOVA when comparing statistically significant difference between the means of three or more independent groups that have been split on two variables, a two-way ANOVA using Šidák’s multiple comparisons test was used. All statistical analyses were performed using Prism 6.0 (GraphPad Software) or Excel. Data are presented as the mean ± SD. and All p-values are shown or ns for non-significant p-value (p > 0.05).

## Acknowledgments

We would like to thank Dr. Vanda Lennon for her feedback, Dr. Emma Goddery and Dr. Roman Khadka for discussions and Ryan Meloche for assisting with logistics of dual MRI. Some graphics were prepared on Biorender.com. Tetramers were provided by the NIH tetramer facility. **Funding sources:** R01NS103212 (AJJ), RF1NS122174 (AJJ), and F31NS116924 (CEF), R01AI167847 (JTH), Nanostring-Glial Panel Grant (CEF).

## Supplementary Table Figure Legends

**Supplementary Table 1.** IPA Causal network analysis identifies MHC Class I family as the number one master regulator of experimental cerebral malaria pathology. Causal network analysis was performed using the Ingenuity Pathway Analysis application. Table includes normalized data from causal network analysis which identifies master regulators of the most highly enriched signaling pathways in the Nanostring dataset comparing uninfected C57BL/6 brain vs. C57BL/6 brains with ECM. Master regulators of causal networks are listed in order of significance, with MHC Class I family identified as most significant causal network of experimental cerebral malaria pathology. P-value of overlap was adjusted using the Benjamini-Hochberg correction method.

**Supplementary Table 2.** Differentially expressed genes in human endothelial cells after *Plasmodium* infection exposure. Using GEO2R analysis platform MHC Class I was identified as one of the most significantly differentially expressed genes and was in the top 5 genes with significant upregulation on HBMECs after *Plasmodium* infection. Listed in order of significance.

**Supplementary Table 3.** Patterns of neuroinflammation determined by cell specific gene enrichment during ECM determined through Mouse Glial Profiling Panel. Table includes the nCounter Mouse Glial Profiling Panel gene lists, probes and corresponding cell type annotations detailing gene expression patterns and their corresponding neuroinflammatory response types.

---

**Supplementary Figure 1.**
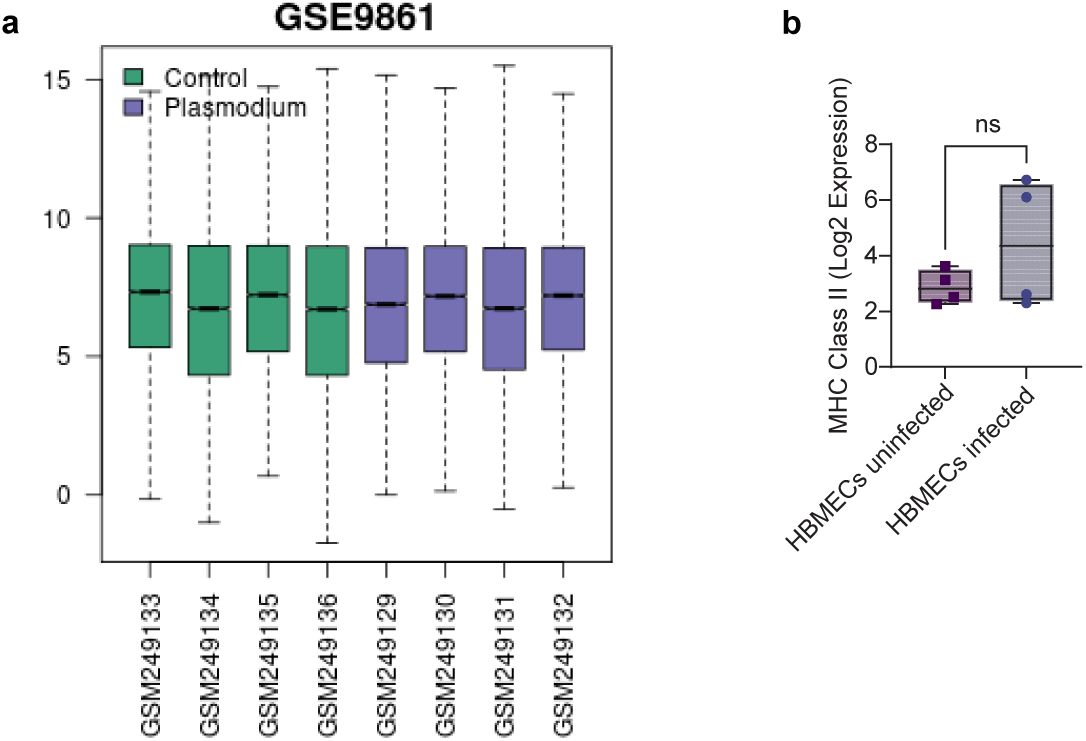
Validation of human data comparison. a) Boxplots show the distribution of the expression values for each human subject. Medians are centered indicating that human datasets are suitable for comparison. Value distribution analysis performed by GE02R Web Application - NCBI GE0. b) MHC Class II Log_2_ expression in control HBMECs vs. Plasmodium exposed HBMECs. Data were analyzed with unpaired t test (b). Box plots present the mean + SD. All p-values are shown or ns for non-significant p-value *(p > 0 05)*.

**Supplementary Figure 2.**
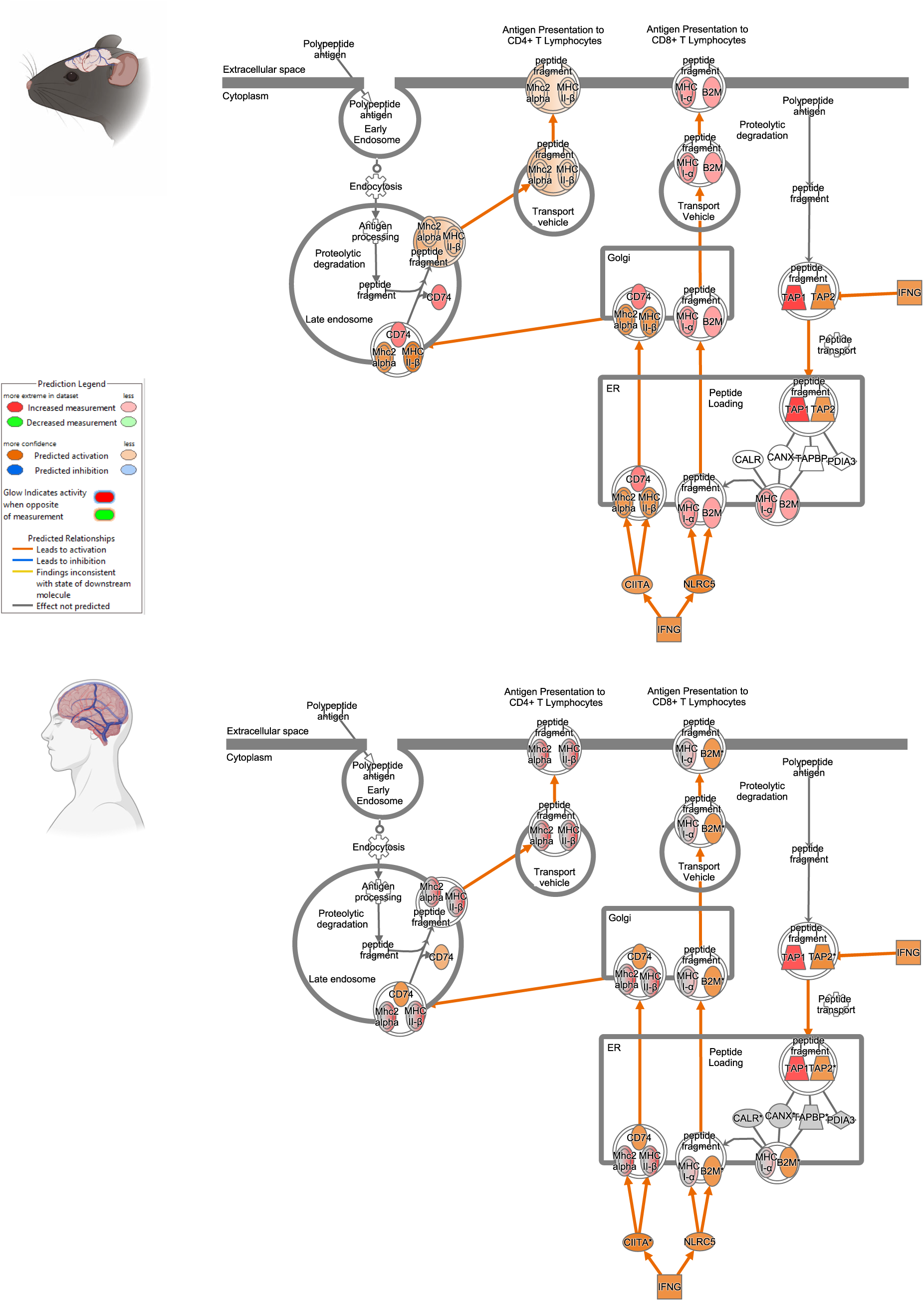
Human and mouse antigen presentation pathways and corresponding genes show similar activation with *Plasmodium* infection. Ingenuity Pathway analyses of mouse and human antigen presentation pathways are depicted in schematics. Coloring of the molecules corresponds to calculated z-scores (algorithm predicting activation or inhibition). Red indicates increased gene activation, orange indicates predicted activation of pathways, green indicates decreased gene activation, blue indicates predicted inhibition of pathways, white indicates a z-score of zero with equal evidence for activation and inhibition, grey indicates that z-score could not be calculated.

**Supplementary Figure 3.**
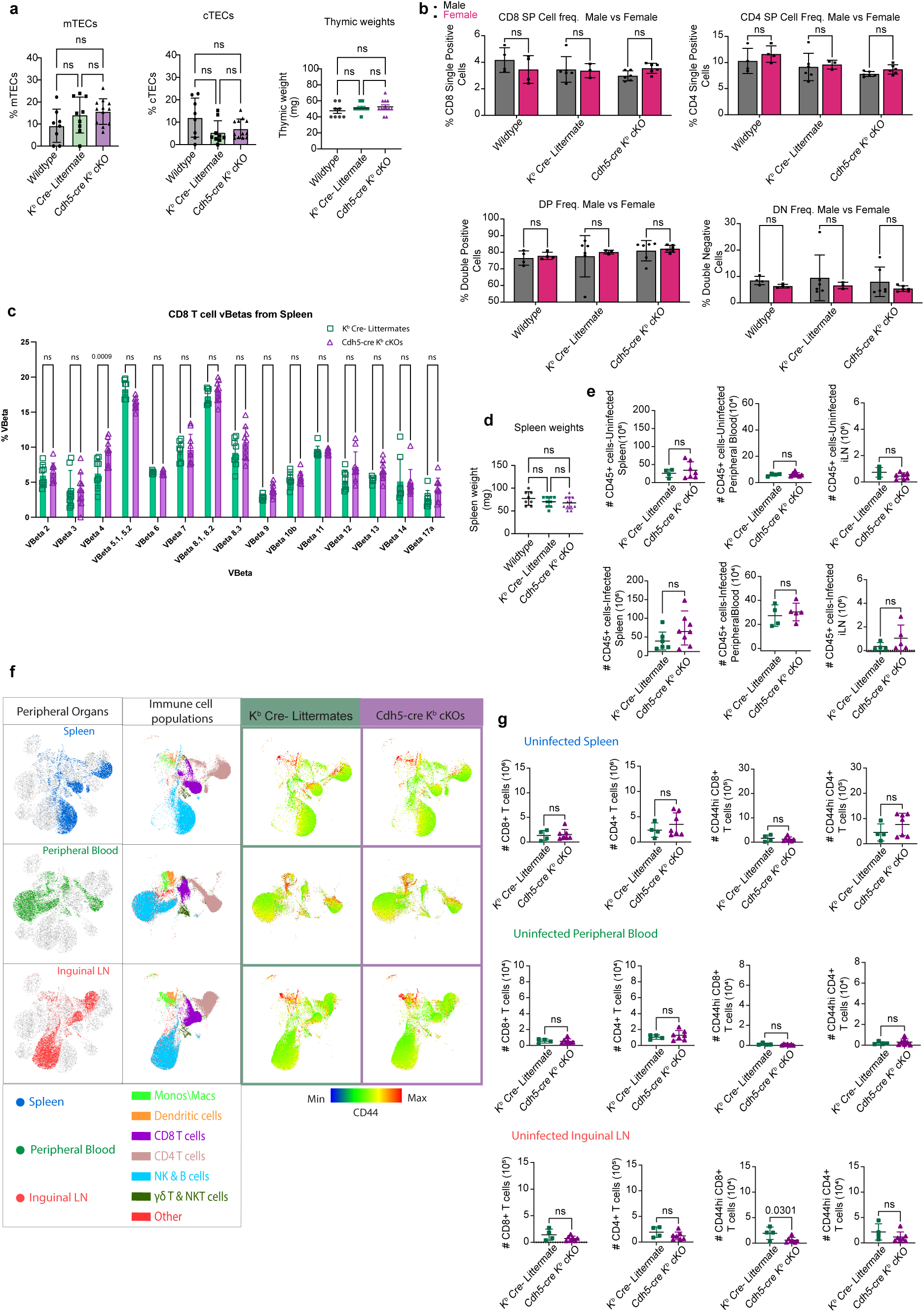
Peripheral immune system is intact K^b^ Cre- littermates and Cdh5-cre K^b^ cKO mice. a) Percentage of mTECs and cTECs quantified from thymi of naïve mice across groups and thymic weights of naïve mice across groups. b) Comparison of percentages of thymic T cell subsets in males vs females across groups in naïve mice. c) Percentages of respective VBeta TCRs out of CD8 T cells of the spleen in naïve mice. d) Splenic weights of naïve mice across groups. e) Absolute counts of CD45+ immune cells in respective peripheral organs of infected and naïve mice. f) Representative UMAPs depicting peripheral organs and immune cell populations and respective heats maps with CD44 activation marker expression. g) The scatter plots show quantified absolute cell counts of CD8 T cells, CD4 T cells and CD44hi CD8 T cells and CD4 T cells in the respective organs depicted in UMAPs. Data were analyzed with unpaired t test, Ordinary 1way ANOVA with multiple comparisons or 2way ANOVA with multiple comparisons. Scatterplots and bar plots present the mean ± SD. All p-values are shown or ns for non-significant p-value *(p > 0.05)*.

**Supplementary Figure 4.**
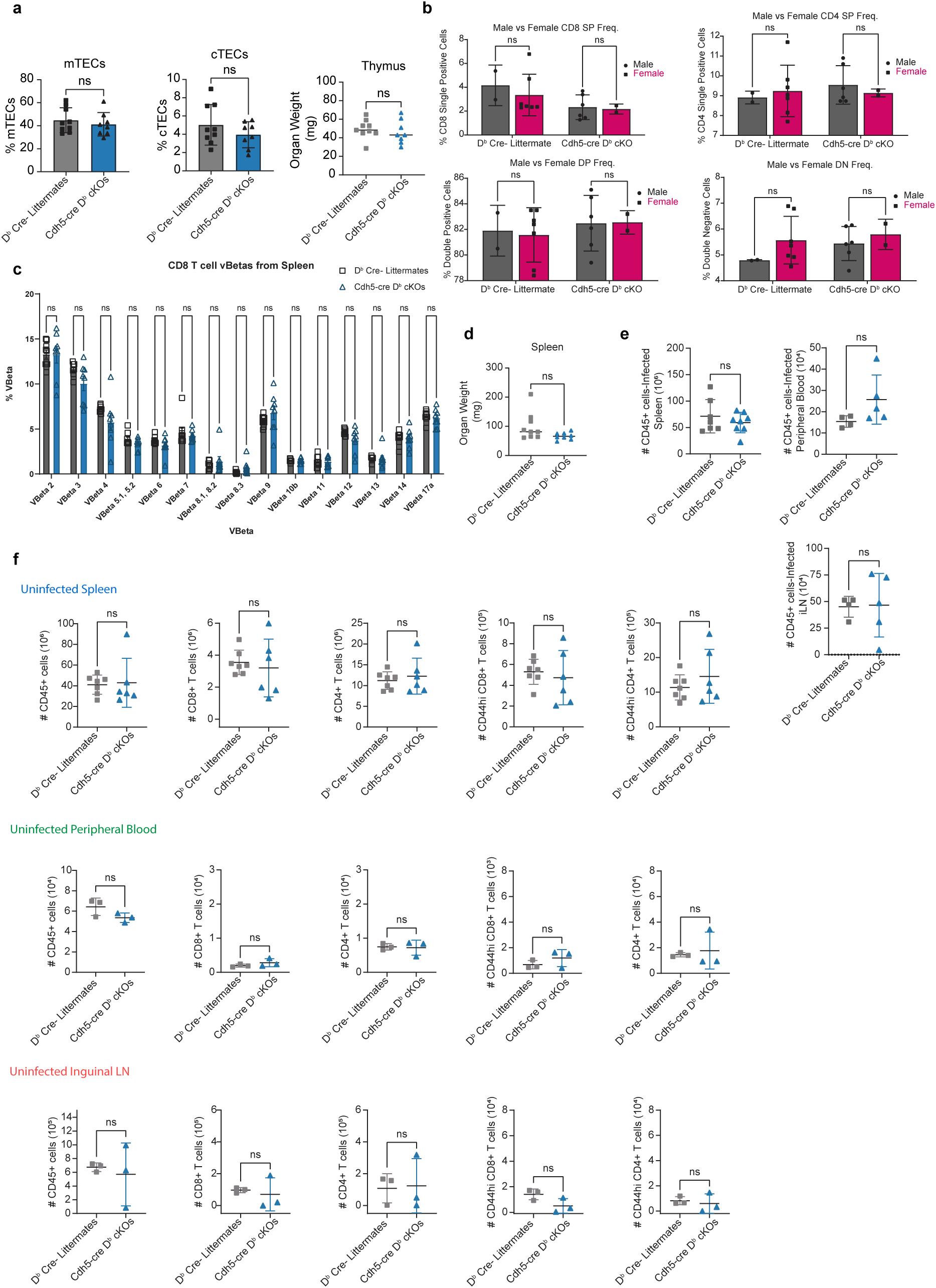
Peripheral immune system is intact in D^b^ Cre- littermates and Cdh5-cre D^b^ cKO mice. a) Percentage of mTECs and cTECs quantified from thymi of naïve mice across groups and thymic weights of naïve mice across groups. b) Comparison of percentages of thymic T cell subsets in males vs females across groups in naïve mice. c) Percentages of respective VBeta TCRs out of CD8 T cells of the spleen in naïve mice. d) Splenic weights of naïve mice across groups. e) Absolute counts of CD45+ immune cells in respective peripheral organs of infected mice. f) Scatter plots show quantified absolute cell counts of CD45+ immune cells, CD8 T cells, CD4 T cells and CD44hi CD8 T cells and CD4 T cells in respective organs of naïve mice. Data were analyzed with unpaired t test, Ordinary 1way ANOVA with multiple comparisons or 2way ANOVA with multiple comparisons. Scatterplots and bar plots present the mean ± SD. All p-values are shown or ns for non-significant p-value *(p > 0.05)*.

**Supplementary Figure 5.**
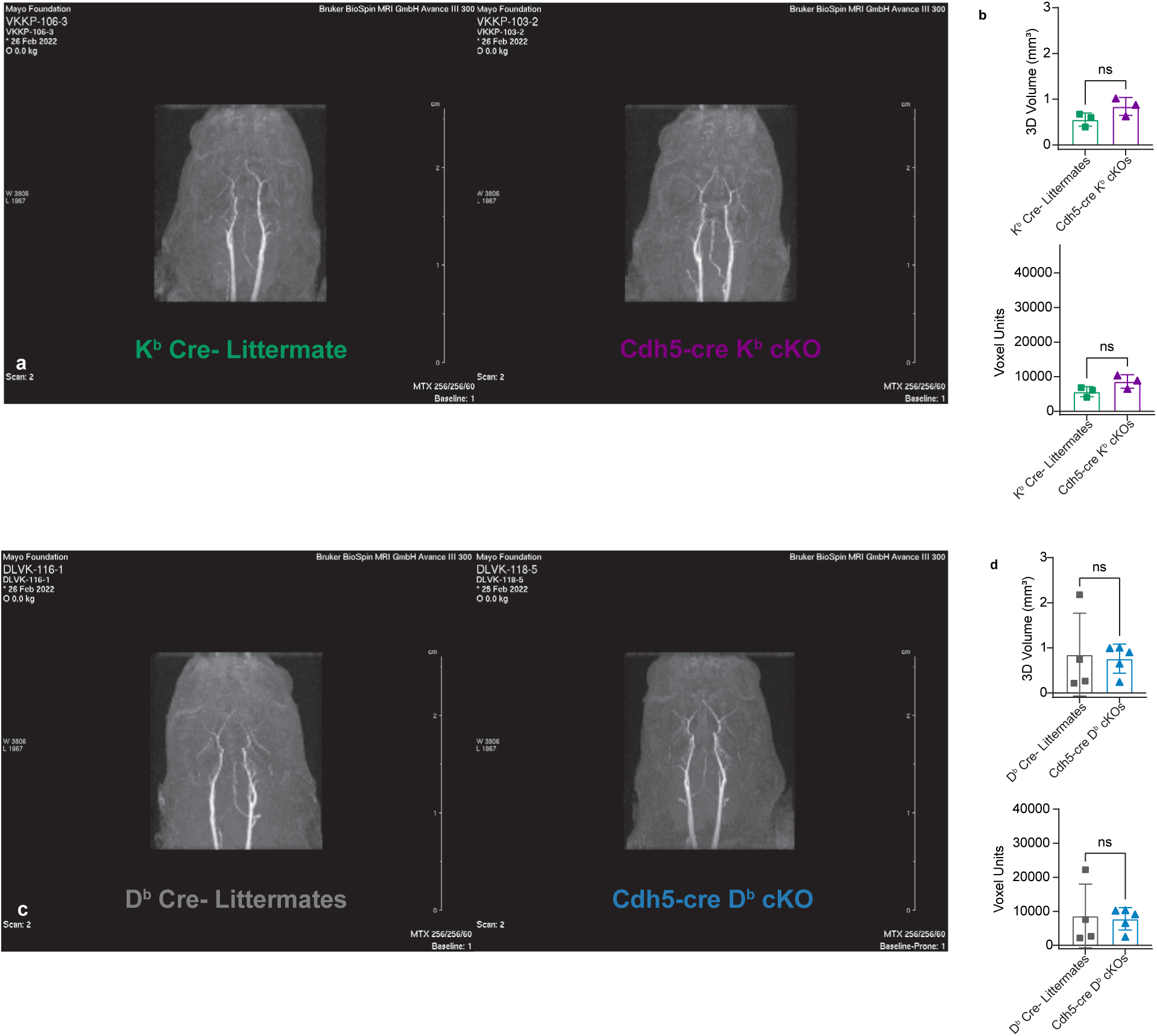
No changes to brain vasculature at baseline in mice with class I deletion on endothelium. a) Representative images from maximum intensity projections (MIPs) of naïve Cdh5-Cre K^b^ cKO and Cre- littermate controls. Mice were IP injected with gadolinium l5 min. before MRIs were performed. b) Quantification of 3D volume of overall brain vasculature enhancement and quantification of 3D pixelated voxel units. c) Representative images from maximum intensity projections (MIPs) of naïve Cdh5-Cre D^b^ cKO and Cre- littermate controls. Mice were IP injected with gadolinium l5 min. before MRIs were performed. d) Quantification of 3D volume of overall brain vasculature enhancement and quantification of 3D pixelated voxel units. Data were analyzed with unpaired t test. Scatterplots present the mean + SD. All p-values are shown or ns for non-significant p-value *(p > 0 05)*.

**Supplementary Figure 6.**
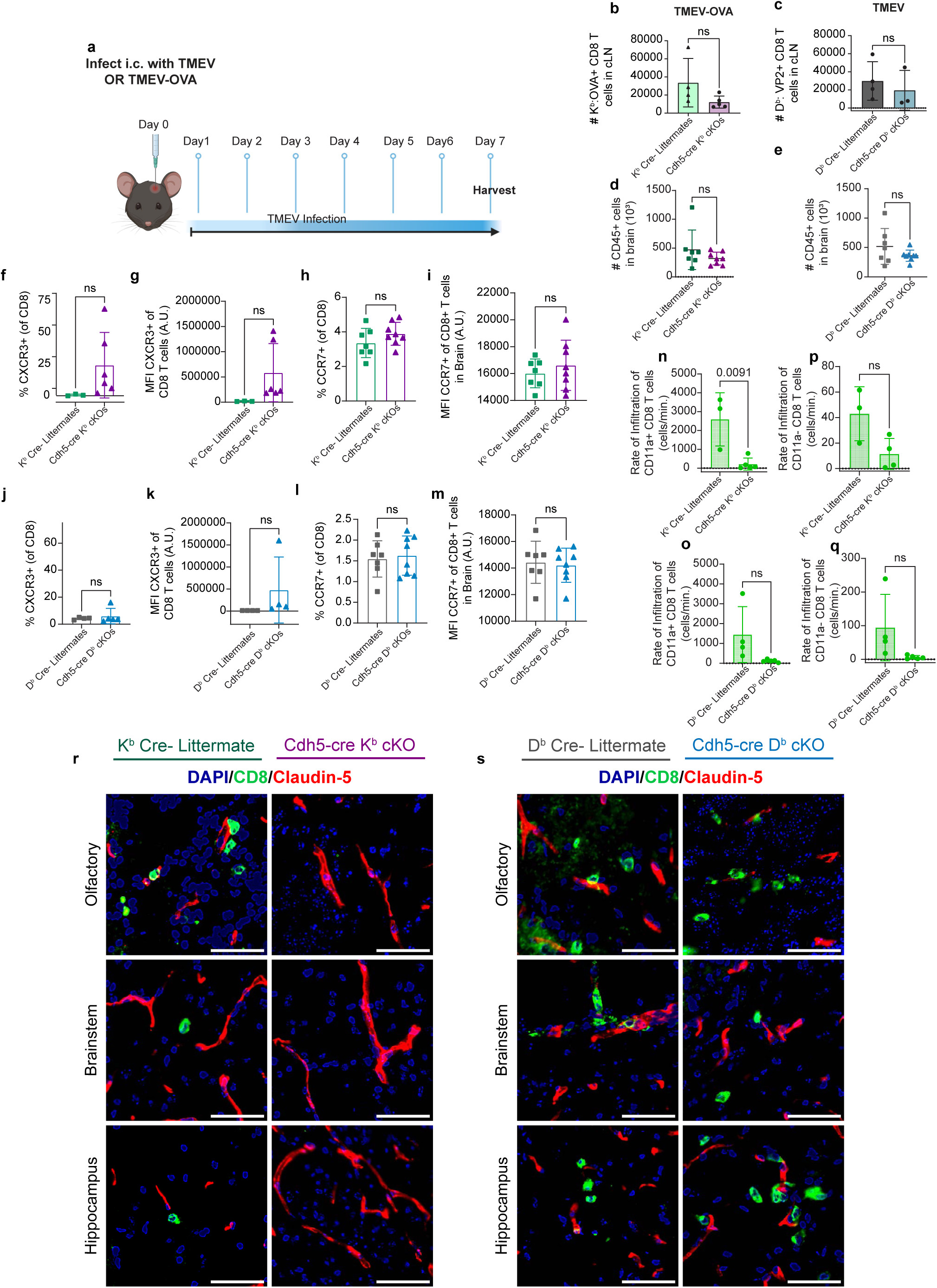
CD8 T cells infiltrate regions of interest except when K^b^ is ablated from vasculature. a) Timeline depicting intracranial infection on Day 0 with TMEV or TMEV-OVA which is allowed to progress until day 7 when brains are harvested for high-dimensional spectral flow cytometry. b) Quantification of absolute counts of K^b^: OVA_257-264_^+^ CD8 T cells in Cdh5-Cre K^b^ cKO and Cre- littermate cLNs day 7 of TMEV-OVA infection. c) Quantification of absolute counts of D^b^: VP2^+^ CD8 T cells in Cdh5-Cre K^b^ cKO and Cre- littermate cLNs day 7 of TMEV infection. d) Absolute counts of CD45+ immune cells in the brains of Cdh5-Cre K^b^ cKO and Cre- littermate brains during ECM. e) Absolute counts of CD45+ immune cells in the brains of Cdh5-Cre D^b^ cKO and Cre- littermate brains during ECM. f) Quantified percentage of CD8s that are CXCR3+ in brains of Cdh5-Cre K^b^ cKO and Cre- littermates during ECM. g) Quantification of median fluorescence intensities of CXCR3 on CD8 T cells in the brains of Cdh5-Cre K^b^ cKO and Cre- littermates during ECM. h) Quantified percentage of CD8s that are CCR7+ in brains during ECM. i) Quantification of median fluorescence intensities of CCR7 on CD8 T cells in the brain during ECM. j) Quantified percentage of CD8s that are CXCR3+ in brains of Cdh5-Cre D^b^ cKO and Cre- littermates during ECM. k) Quantification of median fluorescence intensities of CXCR3 on CD8 T cells in the brains of Cdh5-Cre D^b^ cKO and Cre- littermates during ECM. l) Quantified percentage of CD8s that are CCR7+ in brains during ECM. m) Quantification of median fluorescence intensities of CCR7 on CD8 T cells in the brain during ECM. n) Calculation of rate of infiltration of antigen-experienced CD11a+ CD8 T cells into the brain from IV labeled vasculature, measured in cells/min in infected K^b^ Cre- littermates and Cdh5-Cre K^b^ cKOs. o) Calculation of rate of infiltration of antigen-experienced CD11a+ CD8 T cells into the brain from IV labeled vasculature, measured in cells/min in infected D^b^ Cre- littermates and Cdh5-Cre D^b^ cKOs. p) Calculation of rate of infiltration of CD11a- CD8 T cells that have not encountered cognate peptide:MHC into the brain from IV labeled vasculature, measured in cells/min in infected K^b^ Cre- littermates and Cdh5-Cre K^b^ cKOs. q) Calculation of rate of infiltration of CD11a- CD8 T cells that have not encountered cognate peptide:MHC into the brain from IV labeled vasculature, measured in cells/min in infected D^b^ Cre- littermates and Cdh5-Cre D^b^ cKOs. Confocal imaging was performed. All images were acquired at 20X and zoomed in to 400X using FIJI. r) Representative immunofluorescence images of CD8 T cell infiltration in ROIs affected by ECM showing infiltration into these regions in the K^b^ Cre- littermates but not in the Cdh5-Cre K^b^ cKOs. Scale bar, 50 µM. s) Representative immunofluorescence images of CD8 T cell infiltration in ROIs affected by ECM showing infiltration into these regions in the D^b^ Cre- littermates and in the Cdh5-Cre D^b^ cKOs. Scale bar, 50 µM. Data were analyzed with unpaired t test. Bar plots present the mean ± SD. All p-values are shown or ns for non-significant p-value *(p > 0.05)*.

**Supplementary Figure 7.**
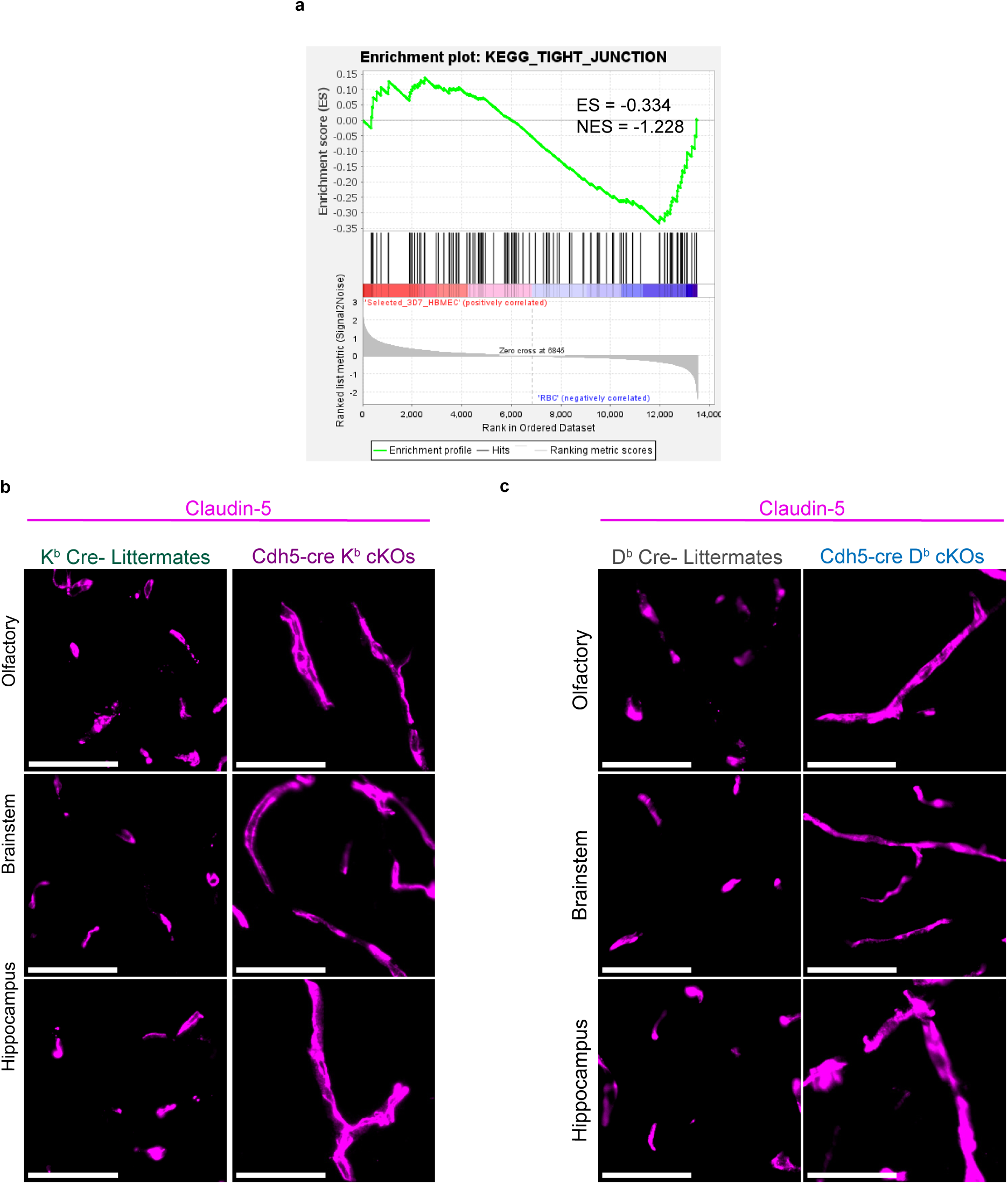
Tight junction protein alterations occur in human and mice with cerebral malaria. GSEA was performed on HBMEC transcriptomic data. a) Enrichment plot showing a negative enrichment (downregulation) of molecular signature for tight junction gene set. The y-axis represents enrichment score (ES) and on the x-axis are genes (vertical black lines) represented in gene sets. The green line connects points of ES and genes. ES is the maximum deviation from zero as calculated for each gene going down the ranked list and represents the degree of over-representation of a gene set at the top or the bottom of the ranked gene list. The colored band at the bottom represents the degree of correlation of genes with the phenotype (red for positive and blue for negative correlation). Significance threshold set at nominal p-value < 0.05. Gene sets for tight junctions were significantly negatively enriched indicating downregulation in HBMECs during infection. Confocal imaging was performed. All images were acquired at 20X and zoomed in to 300X using FIJI. b) Immunofluorescence staining of tight junction protein Claudin-5 in ROI in PbA infected Cdh5-Cre K^b^ cKOs and K^b^ Cre- littermates brains day 6. Scale bar, 50 µM. c) Immunofluorescence staining of tight junction protein Claudin-5 in ROI in PbA infected Cdh5-Cre D^b^ cKOs and D^b^ Cre- littermates day 6. Scale bar, 50 µM.

**Supplementary Figure 8.**
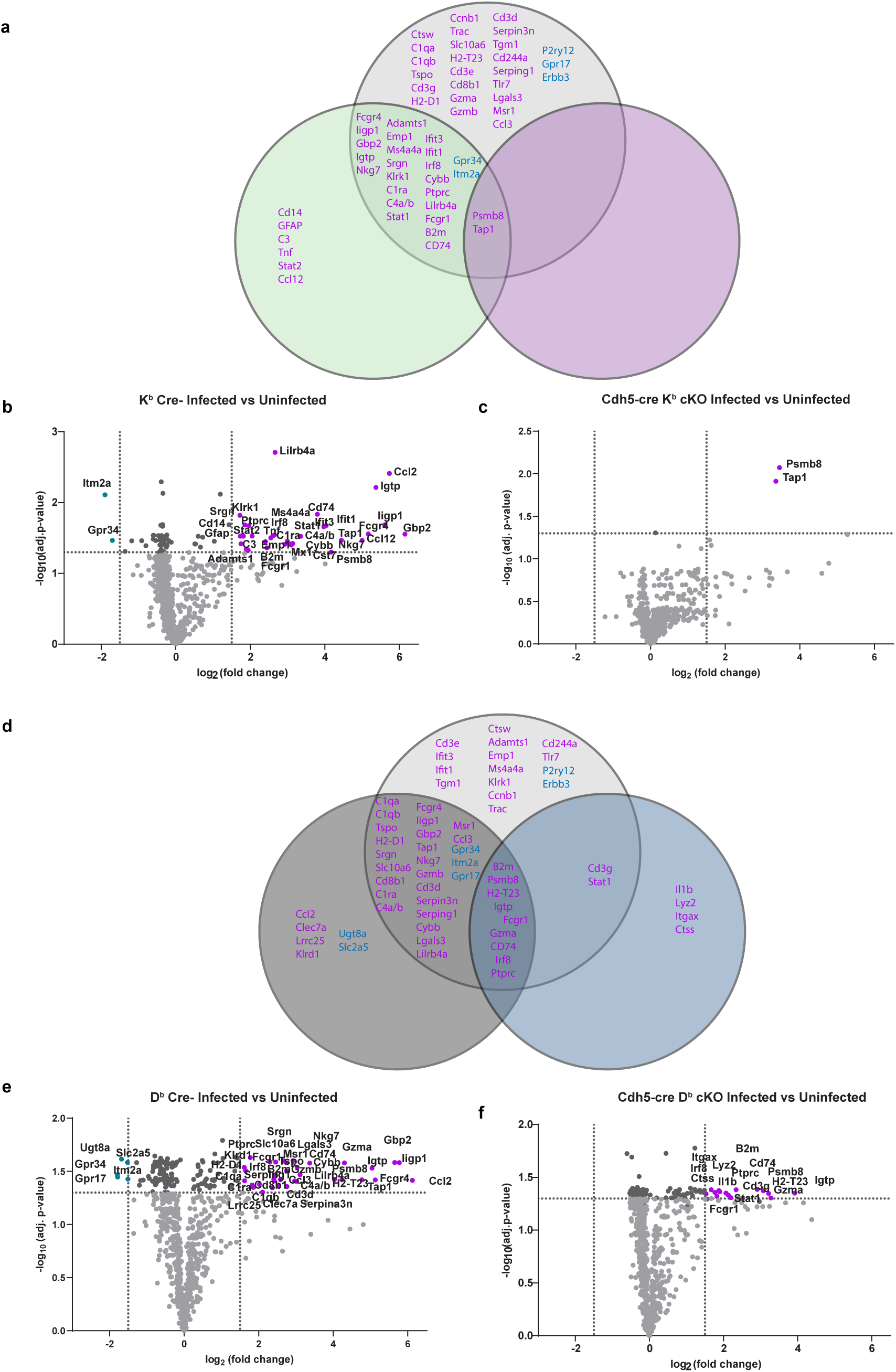
Discrete endothelial class I molecules regulate differential downstream gene expression. a) Venn diagrams of differentially expressed genes are visualized for comparison of differential class I regulation in wildtype, K^b^ Cre- littermates and Cdh5-cre K^b^ cKO mice. Volcano plots of differentially expressed genes in whole brains of uninfected versus PbA infected groups of mice at endpoint during ECM. Significantly upregulated genes are indicated on upper right (purple dots). Significantly downregulated genes are indicated on upper left (turquoise dots). Horizontal dotted line indicates cutoff of adjusted p-values (-Log10 of adjusted p-value 1.3 corresponding to a threshold of p-value of 0.05). Vertical dotted line indicates cutoff of log2 Fold change (-1.5 to 1.5). b) Volcano plots of differentially expressed genes in infected K^b^ Cre- littermates vs Uninfected K^b^ Cre- littermates. c) Volcano plots of differentially expressed genes in infected Cdh5-cre K^b^ cKO mice vs Uninfected Cdh5-cre K^b^ cKO mice. d) Venn diagrams of differentially expressed genes are visualized for comparison of differential class I regulation in wildtype, D^b^ Cre- littermates and Cdh5-cre D^b^ cKO mice. e) Volcano plots of differentially expressed genes in infected D^b^ Cre- littermates vs Uninfected D^b^ Cre- littermates. f) Volcano plots of differentially expressed genes in infected Cdh5-cre D^b^ cKO mice vs Uninfected Cdh5-cre D^b^ cKO mice. P-values of overlap were adjusted using the Benjamini-Hochberg correction method.

